# Quantitative Proteomics Identifies PTP1B as Modulator of B Cell Antigen Receptor Signaling

**DOI:** 10.1101/2021.03.30.437652

**Authors:** Jennifer J. Schwarz, Lorenz Grundmann, Thomas Kokot, Kathrin Kläsener, Sandra Fotteler, David Medgyesi, Maja Köhn, Michael Reth, Bettina Warscheid

## Abstract

B cell antigen receptor (BCR) signaling is initiated by protein kinases and limited by counteracting phosphatases that currently are less well studied in their regulation of BCR signaling. We here used the B cell line Ramos to identify and quantify human B cell signaling components. Specifically, a protein tyrosine phosphatase profiling revealed a high expression of the protein tyrosine phosphatase 1B (PTP1B) in Ramos and human naïve B cells. The loss of PTP1B leads to increased B cell activation. Through substrate trapping in combination with quantitative mass spectrometry, we identified 22 putative substrates or interactors of PTP1B. We validated Igα, CD22, PLCγ1/2, CBL, BCAP and APLP2 as specific substrates of PTP1B in Ramos B cells. The tyrosine kinase BTK and the two adaptor proteins GRB2 and VAV1 were identified as direct binding partners and potential substrates of PTP1B. We showed that PTP1B dephosphorylates the inhibitory receptor protein CD22 at phosphotyrosine 807. We conclude that PTP1B negatively modulates BCR signaling by dephosphorylating distinct phosphotyrosines in B cell specific receptor proteins and various downstream signaling components.

## INTRODUCTION

Protein tyrosine phosphorylation plays a crucial role in the regulation of signal transduction processes and a tight regulation is crucial for cell fate decisions (Hunter 2000). Over the years, knowledge about protein tyrosine kinases (PTKs) has considerably increased, whereas information about protein tyrosine phosphatases (PTPs) and their role in the initiation or termination of signaling is still incomplete. The human genome encodes 109 PTPs, which belong to three different enzyme classes (Damle and Köhn 2019). The vast majority of those belong to class I, which is defined by their signature motif HC(X)_5_R (Li *et al*. 2013). To this class belong the “classical PTPs”, consisting of both receptor type and non-receptor type phosphatases (Alonso *et al*. 2004). In general, the substrate specificity of PTPs is defined by their catalytic domain in combination with domains for localization and substrate recruitment in a cellular context (Tiganis and Bennett 2007).

PTP1B is a classical, ubiquitously expressed non-receptor type phosphatase comprising an N-terminal catalytic domain, two proline-rich motifs and an endoplasmic reticulum (ER)-targeting domain in its C-terminal region (Yip *et al*. 2010). The active site of PTP1B contains the catalytic cysteine residue which is needed for the dephosphorylation of substrate tyrosine residues. The two proline-rich motifs enable the binding of src-homology 3 (SH3) domain-containing proteins, such as p130Cas (Liu *et al*. 1996). Notably, PTP1B is targeted to the ER membrane with a short tail entering the ER lumen (Frangioni *et al*. 1992), whereas both the proline-rich motifs and the catalytic domain are residing in the cytosol. Known substrates of PTP1B include several receptor tyrosine kinases (RTKs) and receptor-associated kinases including the insulin receptor (IR), the epidermal- (EGFR), platelet-derived- (PDGFR), and insulin-like growth factor receptor (IGFR) as well as Janus kinase 2 (JAK2) in various non-hematopoietic cell types (Elchebly *et al*. 1999; Flint *et al*. 1997; Fan *et al*. 2013; Haj *et al*. 2003; Myers *et al*. 2001).

PTP1B is also expressed in the hematopoietic cell linage (Lu *et al*. 2008; Martin-Granados *et al*. 2015; Xu *et al*. 2008). Mice deficient for both, PTP1B and the tumor suppressor p53, have an accumulation of B cells in bone marrow and lymph nodes, leading to an increased susceptibility for B cell lymphomas (Dubé *et al*. 2005). Furthermore, mice with a B cell-specific deletion of PTP1B develop systemic autoimmunity when aged, with elevated anti-dsDNA antibodies in their sera, spontaneous germinal centre formation and glomerulonephritis (Medgyesi *et al*. 2014). Analysis of splenic B cells in these mice revealed p38 MAPK-pY182 as a specific substrate of PTP1B.

In this work, we studied the proteome of the human Burkitt’s lymphoma B cell line Ramos at absolute quantitative scale. By comparing quantitative proteome and transcriptome data and by profiling the expression of classical PTPs in Ramos and naïve B cells, we revealed a role of PTP1B in signal transduction from the B cell antigen receptor (BCR). Loss of PTP1B resulted in increased phosphorylation of several BCR signaling components. Through a quantitative *in vivo* substrate trapping approach, we identified 22 substrates and/or binding partners of PTP1B in human Ramos B cells, many of which having B cell-specific functions. We validated a set of 10 candidates, including the BCR signaling subunit alpha chain (Igα/CD79A) and cluster of differentiation-22 (CD22), an important inhibitory receptor for BCR signaling. Moreover, we identified in the intracellular domain of CD22 the tyrosine dephosphorylated by PTP1B. Thus, we gained new molecular insight how PTP1B controls BCR signaling and, thereby, protects B cells from hyperactivity and auto-reactivity.

## RESULTS

### The Human Ramos B Cell Proteome at Absolute Quantitative Scale

To explore the human B cell proteome, we performed a global quantitative MS-based proteomic study of the Burkitt’s lymphoma cell line Ramos used as B cell model system. A label-free MS analysis allowed the estimation of copy numbers of 8,086 proteins applying the proteomic ruler method (Wiśniewski *et al*. 2014) (**Figures 1A, S1A** and **S1B**; **Table S1**). Copy numbers span seven orders of magnitude, varying from proteins such as the dedicator of cytokinesis protein 1 (DOCK10) with only 12 to the histone H4 with 1.13×10^8^ copies per Ramos B cell (**Figure 1A**; **Table S1**). The large majority of proteins (80%) exhibited copy numbers between 1,000 and 1,000,000 and 50% between 6,500 to 165,000 copies per cell. Approximately 30% of the total protein copy number (3.28 x 10^9^) in Ramos B cells derived from only 50 different proteins, many of which are involved in central processes including transcription, translation and energy metabolism (**Figure S1C**). The lymphoma character of Ramos B cells was reflected by the abundance of cancer-associated proteins including the heat shock protein 90, the cytokine macrophage migration inhibitory factor MIF, glutathione S-transferase P and peroxiredoxin-1 (Schulz *et al*. 2012; Tew *et al*. 2011; Nicolussi *et al*. 2017). Among the proteins with B cell-specific functions, the BCR lambda light chain variable region 2-14 exhibited highest abundance (**Table S1**).

**Figure 1:**
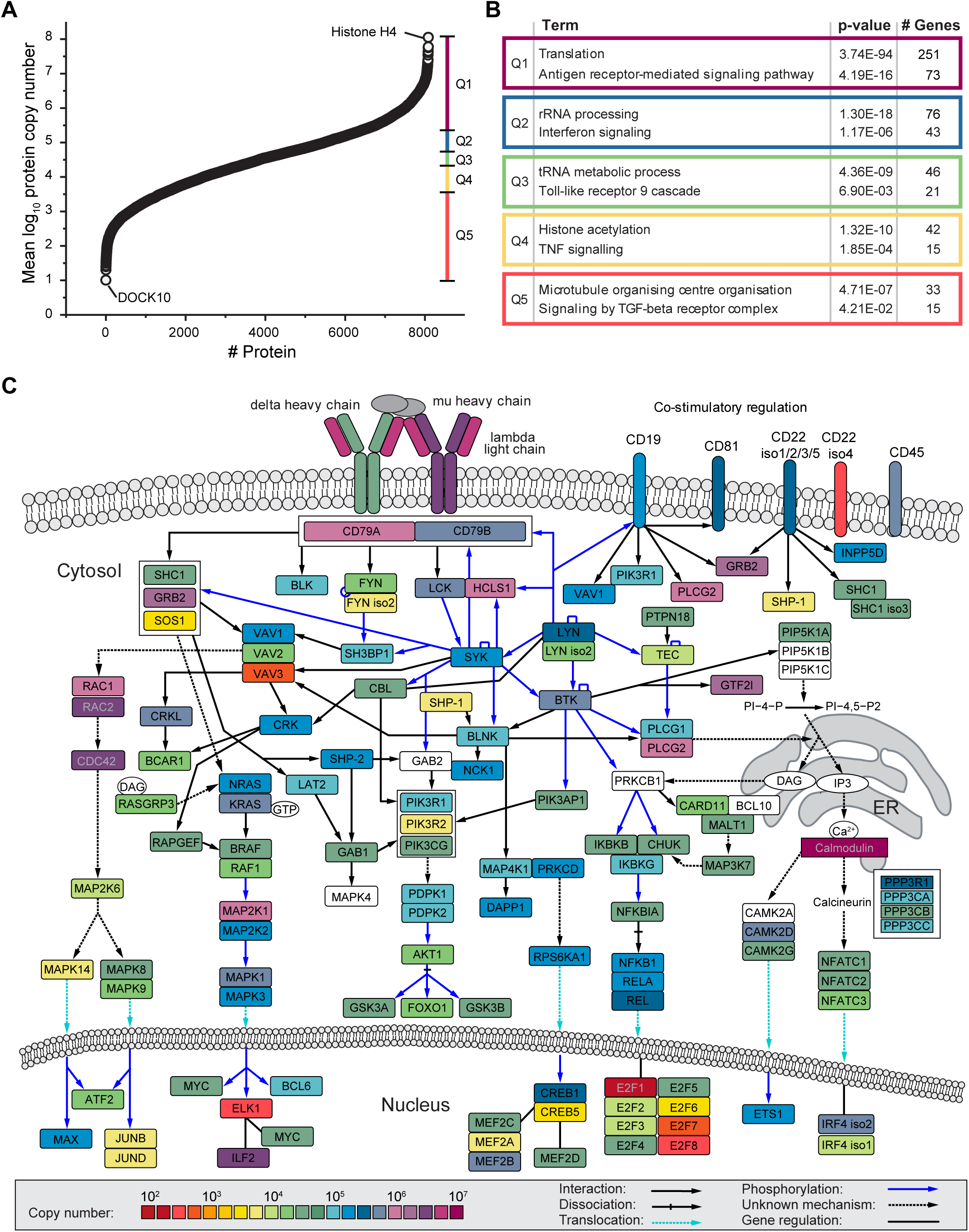
Global absolute quantification of protein expression in Ramos B cells. **A.** Ramos B cell protein copy number plot. Shown are the mean log_10_-transformed copy number values of 8,086 proteins. The dataset was divided into five equal sized quantiles (Q1-Q5) according to copy number rank. **B.** Gene ontology (GO) enrichment analysis of proteins in Q1-Q5, as shown in (A), against the human proteome. For each quantile, significantly enriched processes of the GO term “biological process” and the Reactome pathway are shown with Bonferroni step-down corrected p-values (n=3) and the number of associated genes. **C.** BCR downstream signaling network. The mean copy numbers of individual proteins per Ramos B cell (n=3) are indicated by their color. The pathway was adapted from (Satpathy *et al*. 2015). See also Figure S1 and Supplemental Table S1.

For functional data analysis, we divided the copy number dataset into five quantiles (Q1-Q5) and performed a GO term and Reactome pathway enrichment analysis. In addition to central cellular processes such as translation and transcription, we found in each quantile proteins of distinct signaling pathways to be overrepresented (**Figure 1B**). Among the proteins of highest abundance (Q1), the Reactome pathway term “antigen receptor-mediated signaling pathway” was significantly enriched, highlighting the importance of BCR signaling for Ramos B cell identity, maintenance and function (**Figure 1B**; **Table S2**).

Based on our copy number data and the reported BCR signalosome network (Satpathy *et al*. 2015), we established an absolute quantitative map of the BCR signaling protein network (**Figure 1C**). The IgM class BCR (lambda light chain and mu heavy chain) and the calcium-binding protein calmodulin were most abundant followed by, for example, phospholipase C-gamma-2 (PLCγ2), a crucial component for the initiation of Ca^2+^ signaling. Interestingly, the B cell co-receptors CD22, CD19 and CD81 were considerably less abundant than the IgM-BCR. Other abundant proteins were the hematopoietic lineage cell-specific protein HCLS1, which becomes tyrosine phosphorylated on multiple sites upon BCR stimulation (Yamanashi *et al*. 1993), and the growth factor receptor-bound protein 2 (GRB2), an central adaptor protein which couples several membrane receptors to Ras signaling (Lowenstein *et al*. 1992). In comparison to GRB2, the Ras guanine exchange factor son of sevenless 1 (SOS1) was found to be three orders of magnitude lower in abundance and, thus, it may represent a signaling-limiting factor in B cells. Among the identified transcription factors, interleukin enhancer-binding factor 2 (ILF2) was most abundant, pointing to an important role of this factor in modulating B cell signaling responses.

### The Abundance Profiles of Protein Tyrosine Kinases and Protein Tyrosine Phosphatases

Protein tyrosine phosphorylation mediated by PTKs such as the spleen tyrosine kinase (SYK), the Src family kinases (SFKs) and the Bruton tyrosine kinase (BTK), is essential for BCR activation and downstream signaling (Petro *et al*. 2000; Aoki *et al*. 1994; Weers *et al*. 1994). Based on our Ramos B cell proteome data, we established the copy number profile of 28 PTKs (**Figure 2A**, **Tables S1**). The C-terminal SRC kinase CSK (617,000 copies), BTK (363,000 copies) and two SFKs, namely LYN isoform 1 (304,000 copies) and lymphocyte kinase LCK (338,000 copies) were most abundant followed by PTK2B (163,000 copies) and SYK (123,000 copies), whereas most other PTKs were considerably lower in abundance with copy numbers between 16,000 and 600 per cell. CSK negatively regulates SFKs by the phosphorylation of a carboxy-terminal tyrosine and thereby limits BCR signaling (Hata *et al*. 1994; Chong *et al*. 2005). Interestingly, CSK was expressed at the same level as LYN isoform 1 and LCK together, suggesting that the expression levels of these kinases are precisely regulated.

**Figure 2:**
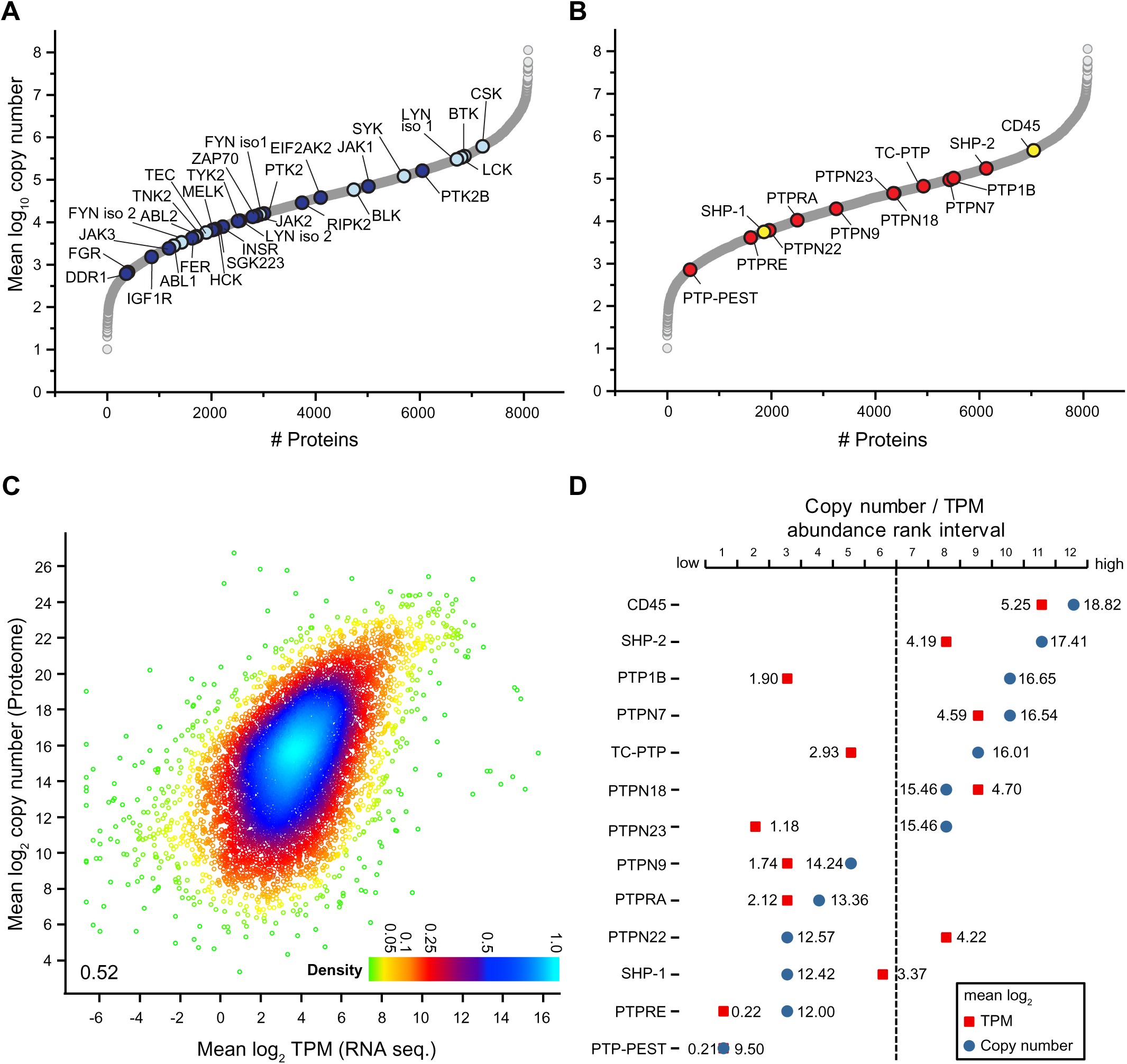
Evaluation of PTKs and classical PTPs in Ramos B cells at quantitative scale. **A.** Copy number profile of 28 different PTKs identified and quantified in Ramos B cells. Src family kinases and other PTKs known to be involved in BCR signaling are shown in light blue, all others in dark blue. **B.** Copy number profile of the 13 classical PTPs identified and quantified in Ramos B cells. CD45 and SHP-1, known to be involved in BCR signaling are highlighted in yellow, the other classical PTPs in red. **C.** Density plot comparing mean log_2_-transformed protein copy numbers of the Ramos B cell proteome analysis against TPM values of the RNA sequencing dataset. A Pearson correlation coefficient of 0.52 was calculated. TPM, transcripts per million **D.** Comparison of the abundance of classical PTPs shown in (B) according to protein (mean log_10_-transformed copy numbers, blue) and mRNA levels (mean log_2_ TPM values, red) of 7,944 proteins and transcripts, respectively. Mass spectrometric and RNA sequencing datasets were divided into 12 equal-sized abundance rank intervals, with the least and most abundant 662 proteins/transcripts in interval 1 and 12, respectively.

In our study of the Ramos B cell proteome, we identified 37 PTPs of which 14 were “classical” PTPs (**Figures 2B** and **S2A**; **Table S1**). From the classical PTPs, the Protein Tyrosine Phosphatase Receptor Type C (PTPRC, also known as CD45) and the src homology region 2 domain-containing phosphatase-1 (SHP-1) play a pivotal role in the control of BCR signaling (Zhu *et al*. 2008; Dustin *et al*. 1999; Cyster and Goodnow 1995; Cornall *et al*. 1998). Most abundant in the group of classical PTPs was CD45 (464,000 copies) with an approximately 1.5-fold higher expression than its substrate Lyn (Katagiri *et al*. 1999). Interestingly, the ubiquitous phosphatase PTP1B (104,000 copies) was ranked third after SHP-2 and expressed at the same level as the kinase SYK, which is essential for BCR downstream signaling (**Figures 2A** and **2B**).

A comparison of the Ramos B cell proteome with the proteome of human naïve B cells (Rieckmann *et al*. 2017) revealed a high correlation (r = 0.70) of protein abundance, including comparable levels for PTKs and classical PTPs (**Figures S2B-S2D**; **Table S3**). The only exception was SHP-1, which was more than 280-fold lower expressed in Ramos (5,600 copies) compared to naïve B cells, whereas CD45 and PTP1B were equally abundant in the two studied B cell populations (**Figures S2C** and **S2D**). Our findings are in line with previous work showing that SHP-1 levels are reduced to 5% in EBV-negative Burkitt lymphomas compared to normal B cells (Delibrias *et al*. 1997).

We next compared protein and mRNA expression levels of Ramos B cells. Mean log_2_ transcript per Million (TPM) values of all protein-coding transcripts (13,611 transcripts) were calculated from RNA sequencing data of Ramos B cells (Qian *et al*. 2014) (**Figure S2E**; **Table S4**). The majority of protein-coding genes followed a normal distribution with a maximum at a log_2_ mean TPM value of approximately 4. Low abundant transcripts caused a divergence from this distribution, but these likely represent lowly expressed genes producing non-functional transcripts (Hebenstreit *et al*. 2011). We thus excluded stochastically expressed transcripts (mean log_2_ TPM value ˂ 0). The GO term enrichment analysis confirmed that many transcripts refer to functions in other cell types (**Table S5**). Mapping of the functional transcripts (9,905; mean log_2_ TPM value ≥ 0) to our proteome data resulted in an overlap of 7,944 proteins with a moderate correlation between TPM and copy number values (r = 0.52) (**Figures 2C** and **2F**; **Table S6)**. We compared the expression levels of 13 classical PTPs quantified at the protein and transcript level (**Tables S1** and **S7**). We found considerable differences between TPM and copy number values and, in most cases, the examined PTPs were more abundant at the protein level (**Figure 2D**). Only SHP-1, PTPN22 and PTPN18 showed higher transcript levels, which points to posttranscriptional or -translational mechanisms that negatively regulate their protein levels. Notably, PTP1B showed the largest discrepancy between transcript and protein levels in B cells. Due to its high abundance at the protein level, a role of PTP1B in B cell signaling can be anticipated.

### Loss of PTP1B Leads to Altered Protein Phosphorylation upon BCR stimulation

To investigate the function of PTP1B in Ramos B cells, we generated PTP1B knockout (KO) cells using the CRISPR-Cas9 technology (Le Cong *et al*. 2013). The PTP1B deficiency was confirmed by immunoblot and MS analysis (**Figures S3A** and **S3B**) and quantitative proteomic analysis showed that loss of PTP1B did not result in global changes in the B cell proteome (**Figure S3C, Table S8**).

To reveal the impact of PTP1B on B cell signaling, changes in tyrosine phosphorylation levels of downstream signaling factors following BCR activation with anti-λ light chain antibody were monitored for different time points (**Figure 3A**). After 3 and 10 min of BCR stimulation, phosphorylation of SYK at Y525/526 was significantly increased in PTP1B KO compared to wildtype (WT) Ramos B cells (**Figures 3A** and **3B**). In addition, ERK-pT202/Y204 levels were significantly increased in PTP1B KO cells (**Figures 3A** and **3C**).

**Figure 3:**
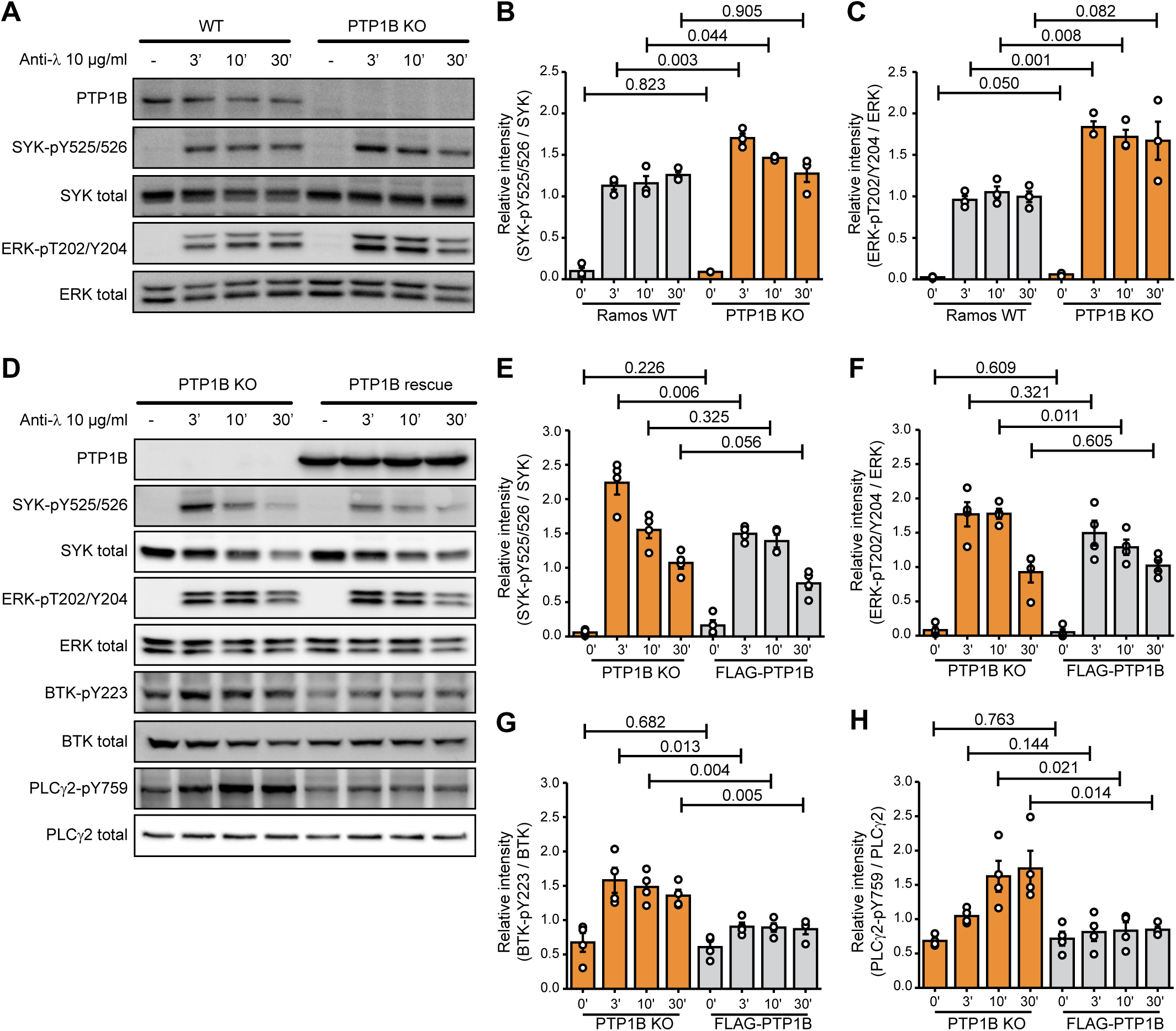
Loss of PTP1B leads to an increase in phosphorylation of SYK-Y525/526, ERK-T202/Y204, PLCγ2-Y759 and BTK-Y223 in Ramos B cells. **A.** Immunoblot analysis of SYK-Y525/526 and ERK-T202/Y204 phosphorylation in PTP1B WT and PTP1B KO Ramos cells stimulated with anti-λ for the indicated time points. Representative immunoblots of three independent biological replicates are shown. **B., C.** Quantification of immunoblot data of SYK-pY525/526 (B) and ERK-pT202/Y204 (C) shown in (A). Signal intensities were normalized to the respective total protein signals and a student’s t-test was performed (n=3). Error bars represent the SEM. **D.** Immunoblot analysis of PTP1B KO and FLAG-PTP1B induced (PTP1B rescue) Ramos cells stimulated with anti-λ for the indicated time points. Representative immunoblots of three independent biological replicates are shown. **E.-H.** Quantification of immunoblot data of SYK-pY525/526 (E), ERK-pT202/Y204 (F), BTK-pY223 (G) and PLCγ2-pY759 (H) shown in (D). Signal intensities were normalized to the respective total protein signals and a student’s t-test was performed (n=4). Error bars represent the SEM.

To show that changes in phosphorylation depend on PTP1B, we reintroduced FLAG- PTP1B in the KO cells using a two component tetracycline-inducible expression system (Haug *et al*. 2015). After BCR stimulation with anti-λ, the SYK-pY525/526 levels were decreased in the FLAG-PTP1B expressing PTP1B KO cells (**Figures 3D** and **3E**). Similarly, ERK-pT202/Y204, BTK-pY223 and PLCγ2-pY759 levels were increased in PTP1B KO cells and decreased upon FLAG-PTP1B expression (**Figures 3D** and **3F-3H)**.

Taken together, our data point to a regulatory role of PTP1B in BCR signaling by limiting the activation of the downstream signaling components.

To analyze the role of PTP1B in intracellular calcium signaling, we measured calcium mobilization after anti-λ stimulation of the BCR by flow cytometry (**Figure S4**). In Ramos B cells expressing FLAG-PTP1B as compared to PTP1B KO cells a similar strong influx of Ca^2+^ ions from the extracellular space into the cytosol was observed. However, the Ca^2+^ levels return faster to the baseline (i.e. level before anti-λ stimulation) in PTP1B KO cells as compared to FLAG-PTP1B expressing cells (**Figure S4B**), which is most apparent at the 300 s time point (**Figure S4C**). This finding indicates a role of PTP1B in restoring the steep calcium concentration gradient between the cytosol and the plasma membrane after a signaling event.

### Global Analysis of the PTP1B-associated Protein Network in Ramos B Cells

To obtain insight into the molecular mode of action by which PTP1B restricts BCR signaling, we generated a FLAG-tagged PTP1B D181A-Y46F substrate trapping mutant (Boubekeur *et al*. 2011), which stabilizes the enzyme-substrate complex and thus allows the recovery of bound tyrosine-phosphorylated substrates by co-immunoprecipitation. In addition, we generated a truncated version that lacks the carboxy-terminal ER-targeting domain (Δ406-435) and localizes to the cytosol (**Figure S5**) to address the question whether the substrate spectrum of PTP1B is determined by its subcellular localization. With this aim we performed a quantitative substrate trapping study that enables to identify and directly compare substrates and/or binding partners of full-length and truncated PTP1B via SILAC-MS (**Figure 4A**). Proteins with a mean SILAC ratio ≥ 2 and a p-value ˂ 0.05 (n=3) were considered as potential PTP1B substrates and/or binding partners (**Figures 4B** and **4C**, **Table S9**). Interestingly, the 12 proteins bound to the truncated cytosolic form of PTP1B represent only a subset of the 22 candidates identified for full-length PTP1B that harbors the ER-targeting domain (**Figure 4D**). These PTP1B interactors include the adaptor protein GRB2, phospholipase C-gamma-1 (PLCγ1) and Rho GTPase-activating protein 12 (ARHGAP12), which have been earlier identified as PTP1B substrates/interactors in other species or cell types (Mertins *et al*. 2008; Banh *et al*. 2016; Ferrari *et al*. 2011; Liu *et al*. 1996). More than one third of the identified substrate candidates carry a SH3 domain (**Table S9**), suggesting that they bind to the proline-rich region of PTP1B. In comparison, only ∼2% of all proteins in the human proteome carry a SH3 domain according to the SMART database (Letunic *et al*. 2015). Network analysis using the STRING database further allowed a classification of the candidates into three main functional groups associated with the GO terms “meiotic cell cycle regulation”, “cargo recognition for clathrin-mediated endocytosis (CME)” and “BCR signaling” to which 10 proteins were assigned (**Figure 4E**; **Table S9**). These data indicate that ER-anchored PTP1B specifically targets a set of proteins with key functions in B cell signaling including the kinase BTK, the inhibitory co-receptor CD22 and the BCR subunits CD79A (Igα) and CD79B (Igβ). Several of the identified PTP1B interactors are involved in calcium signaling and we therefore investigated whether or not the expression of the trapping mutant affects calcium mobilization (**Figure S6**). Indeed, the expression of FLAG-PTP1B D181A-Y46F in PTP1B KO cells strongly reduces the overall calcium flux (**Figures S6B** and **S6D**), indicating that the trapping mutant is capturing a significant pool of substrate candidates that play a role in calcium signaling. Expression of FLAG-PTP1B D181A-Y46F Δ406-435 that lacks the ER-targeting domain also interferes with calcium signaling, but has a milder effect as compared to the full-length trapping mutant (**Figures S6C** and **S6D**).

**Figure 4:**
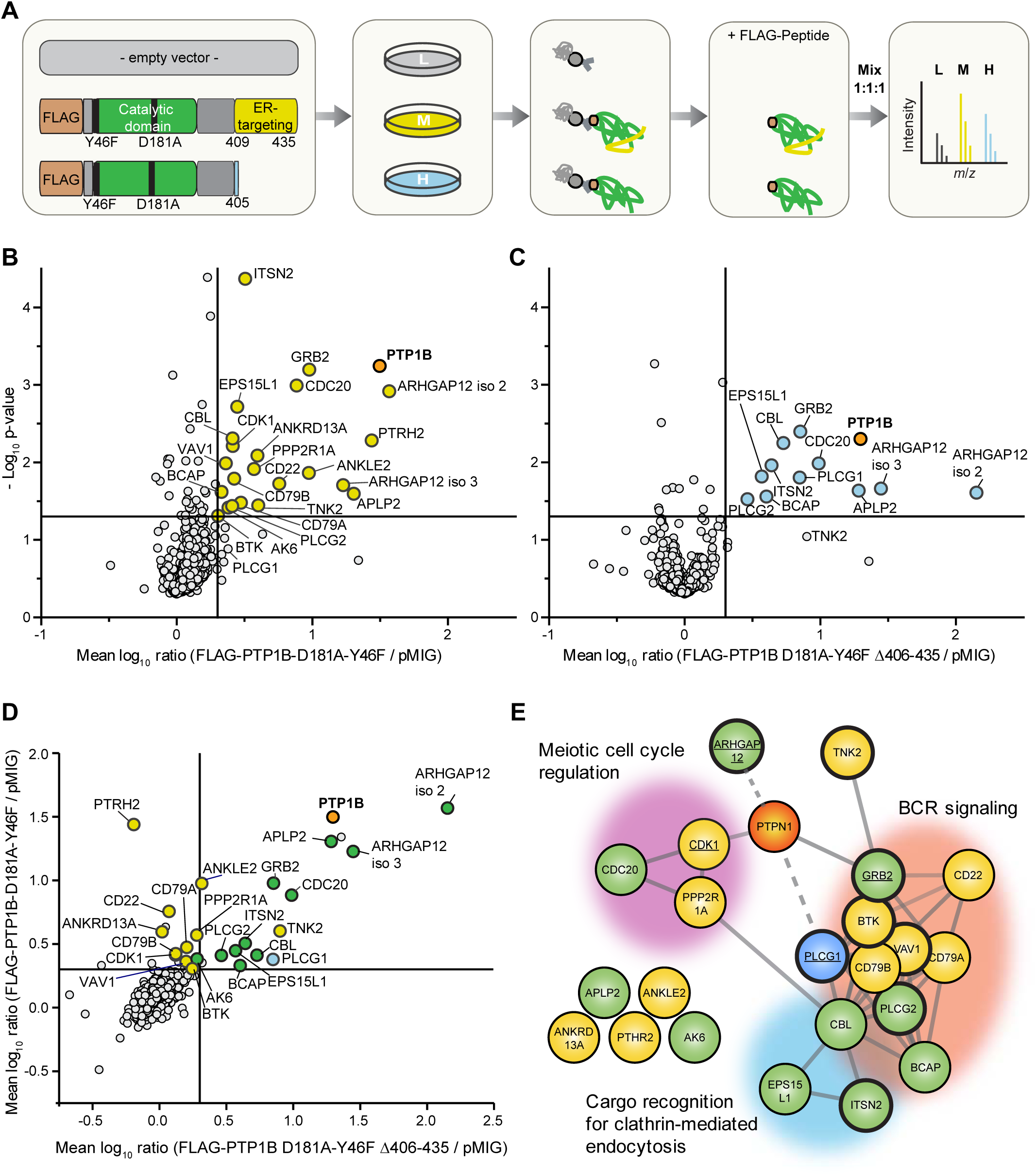
Identification of putative PTP1B substrates in Ramos B cells. **A.** Experimental design. Ramos PTP1B KO cells stably expressing the empty vector control, the FLAG-tagged PTP1B-D181A-Y46F trapping mutant or the FLAG-tagged truncated trapping mutant without the ER-targeting domain were subjected to stable isotope labelling by amino acids in cell culture (SILAC) using “light” (L), “medium-heavy” (M) and “heavy” (H) versions of arginine and lysine. Following SILAC, co-immunoprecipitations using anti-FLAG beads were performed. Bound proteins were eluted by competition with FLAG peptide and eluates were mixed in equal ratio. SILAC samples were separated by SDS-PAGE followed by in-gel digestion of proteins using trypsin and quantitative LC-MS analysis. **B., C.** Scatter plot of proteins eluted with the trapping mutant FLAG-PTP1B-D181A-Y46F (B) and the truncated form of FLAG-PTP1B-D181A-Y46F lacking amino acid residues Δ406-435 (C). Mean log_10_-transformed SILAC ratios were plotted against -log_10_ p-values. Proteins with a minimum fold-change of 2 and a p-value < 0.05 (n=3; right-sided student’s t-test) are highlighted in yellow (B) or in blue (C). The bait PTP1B is shown in orange. **D.** Scatterplot of proteins eluted with both trapping mutants. Mean log_10_-transformed SILAC ratios of proteins in (B) and (C) were plotted against each other. Proteins considered significantly enriched (minimum fold-change of 2 and a p-value < 0.05) with both trapping mutants are marked in green, with FLAG-PTP1B-D181A-Y46F or FLAG-PTP1B-D181A-Y46F Δ406-435 only in yellow and blue, respectively. **E.** STRING database analysis of proteins enriched with the PTP1B substrate trapping mutants. Candidates are grouped into three clusters according to the indicated GO terms. SH3-domain containing proteins are highlighted by bold margins. Known PTP1B (gene name PTPN1) interactors/substrates are underlined.

### Validation of B Cell-specific PTP1B Substrates

For validation of our PTP1B substrate network data, we selected 10 proteins with a focus on proteins involved in BCR signaling and performed anti-FLAG co-immunoprecipitations using PTP1B KO Ramos B cells expressing a FLAG-tagged version of either PTP1B WT or the full-length and truncated form of PTP1B D181A-Y46F (**Figure 5A**). The immunoblot data show that Igα and CD22 exclusively interact with the full-length trapping mutant, which indicates that PTP1B’s localization at the ER is essential for this interaction. In contrast, PLCγ1 and PLCγ2 as well as the B cell adapter for phosphoinositide 3-kinase (BCAP), the amyloid-like protein 2 (APLP2) and the E3 ubiquitin-protein ligase CBL were recovered with both PTP1B trapping mutant versions, with even higher binding to the cytosolic form of PTP1B. Effective binding to the full-length PTP1B trapping mutant was found for BTK, APLP2 and the adaptor proteins GRB2 and guanine nucleotide exchange factor VAV1 all of which also interacted with PTP1B WT to different extent (**Figure 5A**).

**Figure 5:**
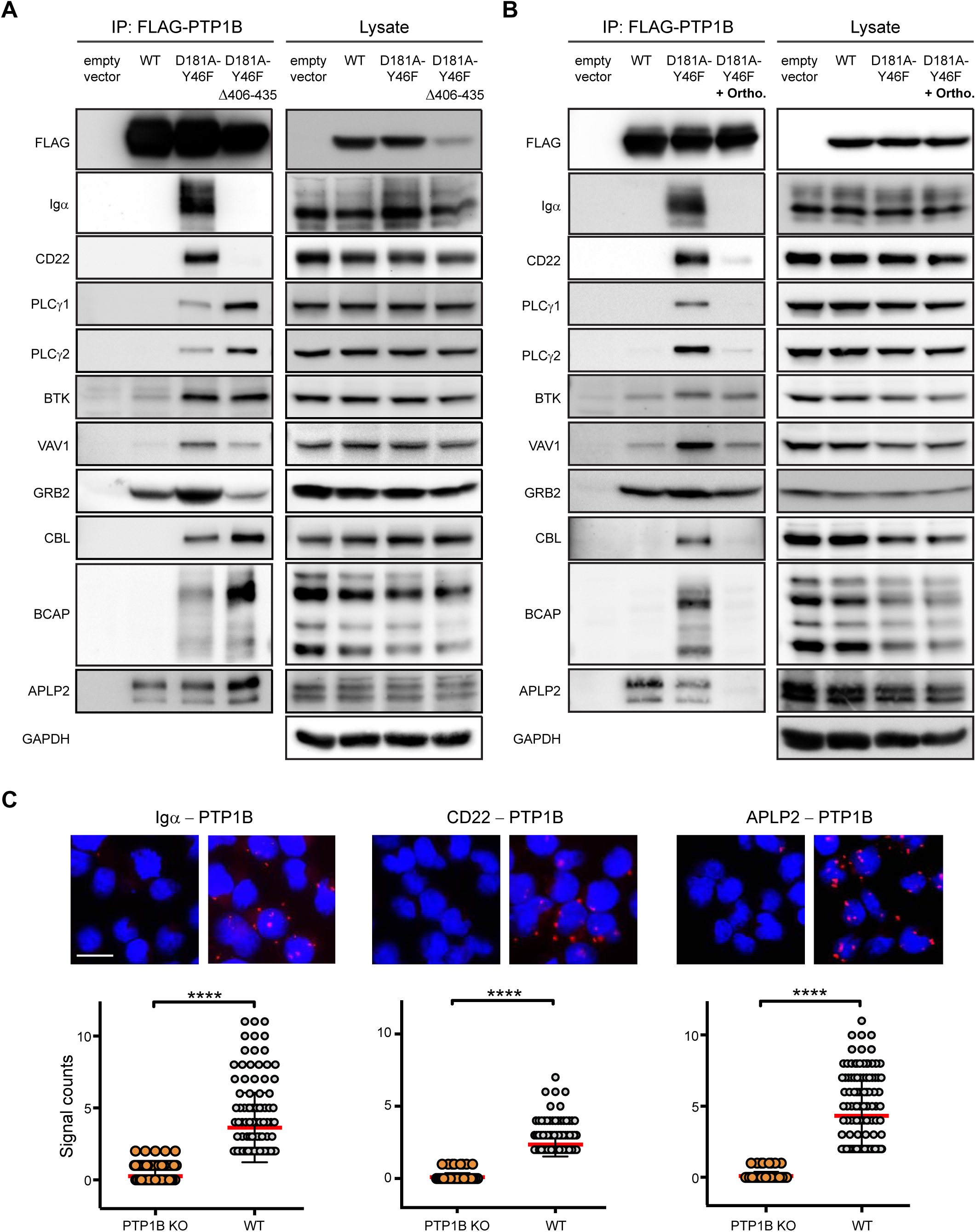
Validation of PTP1B substrates in Ramos B cells. **A, B.** Co-immunoprecipitation experiments were performed using PTP1B KO cells with inducible expression of empty vector control, FLAG-PTP1B WT, FLAG-PTP1B-D181A-Y46F or FLAG-PTP1B-D181A-Y46F Δ406-435 substrate trapping mutants. Eluates were subjected to SDS-PAGE and selected PTP1B substrate candidates (see Fig. 4) were analyzed by immunoblotting (A). To further confirm substrates of PTP1B, cells were lysed in the presence of 1 mM orthovanadate competing for binding and, thus, promoting the release of bound substrate (B). IP, immunoprecipitation; ortho., orthovanadate **C.** PTP1B KO and PTP1B WT Ramos B cells were analyzed for the interaction of PTP1B with endogenous Igα, CD22 or APLP2 by PLA. Data analysis was performed with the CellProfiler software. Each dot represents one cell. Statistical analysis was performed using a Mann-Whitney test (****, p < 0.0001, n=3). Scale bar 10 µm.

To further distinguish between PTP1B binding partners and bound substrates, we used orthovanadate to release the substrate from the catalytic center of PTP1B by binding competition. In the presence of orthovanadate, the interaction with Igα, CD22, PLCγ1/2, CBL and BCAP was virtually abolished, which identifies them as PTP1B substrates (**Figure 5B**). However, the binding of BTK, VAV1 and GRB2 was only reduced, suggesting that this binding is mediated by the SH3 domain of these proteins interacting with the proline-rich sequences of PTP1B.

The interaction of PTP1B with its substrates Igα, CD22 and APLP2 was confirmed by an *in situ* proximity ligation assay (PLA). We observed signals (dots per cell) for PTP1B/Igα, PTP1B/CD22 and PTP1B/APLP2 that were significantly higher in PTP1B WT than PTP1B KO cells (**Figure 5C**), underscoring the specific interaction of these proteins with PTP1B in Ramos B cells.

In summary, we validated Igα, CD22, PLCγ1/2, CBL, BCAP and APLP2 as *in vivo* substrates of PTP1B in B cells. In addition, BTK and the adaptor proteins GRB2 and VAV1 were identified as direct binding partners and potential dephosphorylation targets of PTP1B.

### PTP1B Dephosphorylates Specific Tyrosine Residues in CD22

Among the validated PTP1B substrates was the inhibitory receptor CD22 that counteracts BCR signaling. CD22 carries six tyrosine residues within its intracellular tail sequence, of which three are located within the immunoreceptor tyrosine-based inhibition motifs (ITIMs) of CD22 and one in the ITIM-like motif (**Figure 6A**). All six tyrosine residues are conserved between mouse and human CD22 and with the exception of Y767/Y752, all of them are known to be phosphorylated in vivo (Schulte *et al*. 1992; Yohannan *et al*. 1999; Cornall *et al*. 1998). Following BCR stimulation, GRB2 and SHP-1 are recruited to CD22 (Otipoby *et al*. 2001; Law *et al*. 1996). To identify the tyrosine residue(s) in CD22 that are dephosphorylated by PTP1B, we carried out a peptide-based phosphatase assay. We were able to synthesize four different peptides of human CD22 carrying a central phospho-tyrosine (pY) and sequences from the first ITIM (Y762), the ITIM-like (Y796), the second ITIM (Y822) as well as the GRB2 binding region (Y807). The other two peptides carrying pY752 and pY842 were synthetically not accessible. With these four peptides, we measured the velocity of the dephosphorylation reaction with increasing concentrations of the respective pY-peptides (**Figure 6B**). In comparison to the control peptide KKKKpYPKK (Li and Köhn 2016), all four CD22-pY-peptides were dephosphorylated by recombinant PTP1B. Based on these data, we conclude that these CD22-pY residues are substrates of PTP1B *in vitro*.

**Figure 6:**
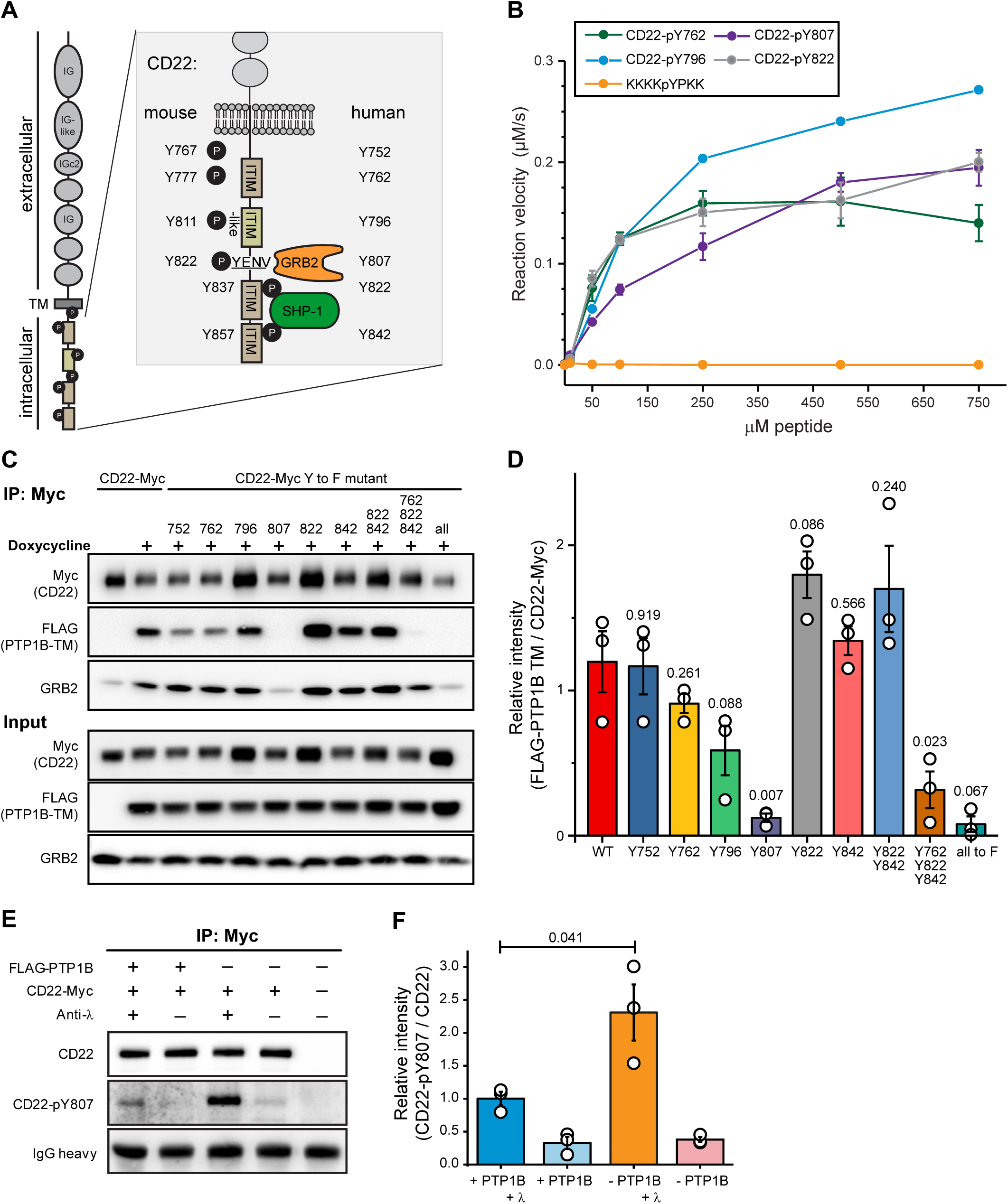
PTP1B dephosphorylates CD22 at pY807. **A.** Schematic representation of the human/mouse CD22 molecule with domain structure and intracellular tyrosine (Y) residues. ITIM, immunoreceptor tyrosine-based inhibitory motif; TM, transmembrane domain; IG, immunoglobulin domain. GRB2, growth factor receptor-bound protein 2; SYK, spleen tyrosine kinase. **B.** Peptide-based phosphatase assay showing the dephosphorylation of four synthetic CD22 peptides by GST-tagged PTP1B. Error bars represent the standard deviation of three independent experiments. **C.** PTP1B KO Ramos cells with inducible FLAG-PTP1B-D181A-Y46F trapping mutant were stably transduced with Myc-tagged CD22 WT or Myc-tagged CD22 site mutants in which single tyrosine residues were replaced by phenylalanine (Y-to-F mutants) as indicated. Following inducible expression of FLAG-PTP1B-D181A-Y46F, co-immunoprecipitations using anti-Myc antibody were performed and bound proteins were eluted. CD22 WT and phosphosite mutants were detected in cell lysates and eluates by SDS-PAGE immunoblotting using anti-Myc antibody. The FLAG-tagged PTP1B trapping mutant was detected using anti-FLAG antibody. Co-immunoprecipitation of GRB2 was detected using a GRB2 antibody. **D.** Quantification of immunoblot data from (C). Anti-FLAG signals (FLAG-PTP1B-D181A-Y46F) were normalized to anti-myc signals (myc-tagged CD22 and site mutants thereof). A student’s t-test was performed testing each mutant against the control and p-values were calculated (n=3). Error bars represent the SEM. **E.** Phosphatase assay in FLAG-PTP1B inducible Ramos cells. Myc-tagged CD22 was immunoprecipitated from untreated or anti-λ stimulated cells, with or without the expression of FLAG PTP1B. Phosphorylation levels of CD22-Y807 were monitored by immunoblot. **F.** Quantification of immunoblots from (E). CD22-pY807 signals were normalized to the signals for immunoprecipitated CD22-myc. A student’s t-test was performed and p-values were calculated (n=3). Error bars represent the SEM.

To examine PTP1B target sites in CD22 in a cellular context, we individually exchanged each of the tyrosine to phenylalanine to prevent phosphorylation. In addition, CD22 mutants were generated in which the tyrosine residues in the second (Y822) and third ITIM (Y842), all ITIMs (Y762, Y822, Y842) or all six tyrosines (6Y-to-F mutant) in its intracellular part were replaced by phenylalanine. Carboxy-terminally Myc-tagged versions of CD22 WT and the different non-phosphorylatable CD22 mutants were stably expressed in PTP1B KO Ramos B cells with inducible expression of the FLAG-PTP1B D181A-Y46F trapping mutant. Since substrate binding is much stronger upon tyrosine phosphorylation, we assumed that tyrosine-to-phenylalanine mutations should reveal the CD22 site(s) targeted by PTP1B. Immunoblot analysis showed that the PTP1B trapping mutant was effectively co-purified with CD22-Myc in Ramos B cells, whereas this interaction was abolished for the 6Y-to-F mutant of CD22 (**Figures 6C** and **6D**). These data confirm that CD22 tyrosine phosphorylation is required for PTP1B binding. The interaction of PTP1B with the CD22 triple mutant was also reduced, which was not the case for the double mutant Y822/842F. The latter was in agreement with the behavior of the single Y822F and Y842F mutants. However, since binding to the single CD22-Y762F mutant was not significantly affected, this site may not be a preferred target site of PTP1B in cells.

In our assay, we found a strongly impaired PTP1B binding for the CD22-Y807F mutant (**Figure 6D**). Our data therefore point to CD22-pY807 as the preferred target site of PTP1B. Of note, human CD22-Y807 (see **Figure 6A**), is part of the Y-E/Q-N-Ψ motif (Ψ, hydrophobic amino acid) that when phosphorylated serves as the binding site for the SH2 domain of GRB2 (Kessels *et al*. 2002) (Poe *et al*. 2000; Otipoby *et al*. 2001; Chen *et al*. 2016). Accordingly, GRB2 binding was reduced to background levels in immunoprecipitates of the CD22-Myc Y807F mutant in Ramos B cells (**Figure 6C**).

To further validate CD22-pY807 as a PTP1B substrate, we performed a cellular PTP1B phosphatase assay using our FLAG-PTP1B-inducible Ramos cells (**Figures 6E** and **S5**). Following anti-λ stimulation of the BCR, CD22 is phosphorylated at Y807 and the phosphorylation level is significantly reduced in cells expressing FLAG-PTP1B (**Figure 6F**).

Taken together, we showed that PTP1B dephosphorylates CD22-pY762, -pY796, - pY807 and -pY822 *in vitro* and identified pY807 as the main target site of PTP1B in Ramos B cells.

## DISCUSSION

Mice with a B cell-specific PTP1B gene knockout develop systemic autoimmunity upon ageing, indicating an important regulatory role for PTP1B in B cell activation (Medgyesi *et al*. 2014). In this work, we aimed at a better understanding of the molecular mode of action of PTPB1 in human B cells using quantitative proteomic and biochemical methods. For our systematic analysis, we chose the Burkitt lymphoma cell line Ramos as a model for mature human B cells. Our absolute proteome-wide quantification revealed a high abundance of PTP1B in Ramos B cells consistent with reported data from naïve B cells (Rieckmann *et al*. 2017). The necessity of quantification at the level of proteins was highlighted by comparing protein copy numbers determined in this work and TPM values calculated from RNA sequencing data of human Ramos B cells (Qian *et al*. 2014). This revealed a considerably higher expression of PTP1B on the protein than on the transcript level. We also compared the abundance of other PTPs regulating B cell signaling such as SHP-1 and CD45 (Katagiri *et al*. 1999; Zhu *et al*. 2008; Cornall *et al*. 1998; Pani *et al*. 1995; Cyster and Goodnow 1995; Dustin *et al*. 1999). We found that SHP-1 was significantly more abundant in naïve B cells compared to Ramos B cells, whereas CD45 copy numbers were consistently high in both of these cells.

Loss of PTP1B in Ramos B cells did not affect steady-state protein levels as shown by SILAC-MS analysis. However, loss of PTP1B altered BCR downstream signaling shown by a significant increase in the levels of SYK-pY525/526, ERK-pT202/Y204, BTK-pY223 and PLCγ2-pY759. These data point to a role of PTP1B in counteracting BCR signaling and, thus, B cell activation. PTP1B was previously shown to dephosphorylate SYK at pY526 in isolated chronic lymphocytic leukemia (CLL) cells from patients (Boelens *et al*. 2009).

In a quantitative substrate trapping approach, we identified 22 potential targets of PTP1B in human Ramos B cells with functions in meiotic cell cycle regulation as well as cargo recognition for clathrin-mediated endocytosis and BCR signaling, two functionally interconnected processes. PTP1B has previously been implicated in the regulation of the cell cycle, since it is a target of cyclin-dependent kinase 1 (CDK1) in mitotic cells (O’Donovan *et al*. 2013). The three PTP1B substrate candidates CBL, intersectin-2 (ITSN2) and epidermal growth factor receptor substrate 15-like 1 (EPS15L1) are associated with clathrin-coated pits and were shown to be tyrosine phosphorylated upon antigen stimulation of the BCR (Matsumoto *et al*. 2009; Satpathy *et al*. 2015). In addition, the ubiquitin-binding protein ankyrin repeat domain 13a (ANKRD13A), a protein involved in EGFR endocytosis, was found in BCR signalosomes (Satpathy *et al*. 2015; Tanno *et al*. 2012). In fact, it was reported that PTP1B regulates the endosomal trafficking of receptor tyrosine kinases (RTKs) (Eden *et al*. 2010) and dephosphorylates signal transducing adapter molecule 2 (STAM2), an endosomal protein co-localizing with clathrin and involved in the sorting of RTKs for lysosomal degradation (Stuible *et al*. 2010). Therefore, it appears likely that CBL, EPS15L and ITSN2 are direct dephosphorylation targets of PTP1B.

Almost half of our identified candidates for PTP1B substrates have a central role in B cells. We successfully validated all of them as PTP1B substrates by Co-IP experiments in combination with orthovanadate treatment. However, we did not target Igβ directly as it forms a disulphide-linked heterodimer with Igα (Siegers *et al*. 2006) and, thus, co-purifies with Igα. In addition to the phosphotyrosine-dependent binding of BTK and PLCγ2 to the PTP1B substrate trapping mutant, we also showed that BTK-pY223 and PLCγ2-pY769 levels were increased in PTP1B KO cells after anti-λ stimulation. Furthermore, we identified and validated VAV1 as PTP1B substrate in Ramos B cells. In MEF cells, pY141 of VAV3 was identified as PTP1B-dependent phosphosite (Mertins *et al*. 2008). Since VAV1 and VAV3 exhibit high sequence similarity (∼75%) and redundant functions (Pearce *et al*. 2004), we propose that PTP1B may dephosphorylate VAV1 at pY142 which corresponds to pY141 in VAV3.

In this work, we demonstrated that PTP1B has a multitude of substrates in B cells. Thus, an important question is how PTP1B reaches its substrates in a cellular context? PTP1B has a proline-rich region through which the interaction with SH3 domain-containing proteins is mediated. For example, the PTP1B-dependent dephosphorylation of breast cancer anti-estrogen resistance protein 1 (BCAR1) was shown to depend on the interaction of PTP1B with the SH3 domain of BCAR1 (Liu *et al*. 1996). Indeed, we found that about one third of the identified PTP1B substrates contain at least one SH3-domain by which they potentially interact with PTP1B. Among these proteins was also GRB2, which contains a SH2 domain flanked by two SH3 domains. By Co-IP and immunoblot analysis we showed that GRB2 directly interacts with WT PTP1B. The binding of GRB2 to the PTP1B substrate trapping mutant was enhanced, but reduced upon orthovanadate treatment. This suggests that GRB2 is primarily an interactor and subsidiary a substrate of PTP1B.

Our data further showed that the interaction of PTP1B with Igα and CD22 requires its ER-targeting domain. This suggest that PTP1B plays an important role at the ER where it may prevent uncontrolled phosphorylation and activation of newly produced receptor components before they are assembled into protein complexes or membrane nano-domains that regulate their activity. Interestingly, the PTP1B-GRB2 interaction was also reduced in case of the truncated PTP1B trapping mutant that lost its location at the ER. Thus, GRB2 may connect CD22 to ER membrane-bound PTP1B. We showed that PTP1B dephosphorylates CD22 at pY807 in its intracellular domain. CD22 directly interacts with the adaptor protein GRB2, with GRB2 binding being dependent on CD22-pY807 (Poe *et al*. 2000). Thus, it is tempting to speculate that PTP1B is recruited to CD22 via binding to GRB2. In this complex, PTP1B may also reach and dephosphorylate other substrates such as PLCγ2, SYK and BTK, which are now in close proximity. Dephosphorylation of CD22-pY807 will then provide a switch for releasing GRB2 and, thereby, protecting CD22 and other substrates from further dephosphorylation by PTP1B.

## EXPERIMENTAL PROCEDURES

### Cell Culture and Cell Lines

Ramos B cells were cultivated in Roswell Park Memorial Institute (RPMI) 1640 Medium with stable L-glutamine (PAN Biotech, Aidenbach, Germany) supplemented with 10% heat-inactivated fetal calf serum (FCS) (Biochrom, Berlin, Germany). For SILAC experiments, cells were cultured for at least six days in RPMI 1640 Medium supplemented with 10% dialyzed FCS (Sigma Aldrich), containing either “heavy” lysine (40 g/l; 13C6, 15N2) and arginine (80 g/l; 13C6, 15N4), “medium” lysine (40 g/l; 4,4,5,5−D4) and arginine (80 g/l; 13C6), or “light” lysine (40 g/l) and arginine (80 g/l). To prevent the conversion of arginine to proline, 80 g/l proline was added. All stable isotope-labelled amino acids were purchased from Cambridge Isotope Laboratories, Andover, MA. The “light” versions of the SILAC amino acids were obtained from Sigma. For Inducible expression of FLAG-PTP1B, FLAG-PTP1B D181A Y46F or FLAG-PTP1B D181A-Y46F Δ406-435 (missing 30 amino acids at the C-terminus) PTP1B KO Ramos were transfected with a doxycycline inducible tet-on vector and exposed to 2 μg/ml doxycycline (Calbiochem, Merck, Darmstadt, Germany) 24 to 48 h before the experiment. Platinum E or Phoenix cells were used as packaging cell lines to generate virus particles. Cells were transfected with the corresponding plasmids using polyethylenimine (1 μg/μL [pH 7]; Polysciences) in a 1:3 ratio. The virus supernatants were harvested 48 h and 72 h after transfection. For each plasmid, 2×10^6^ Ramos B cells were infected with 500 µl virus supernatant for 3x 60 min centrifuging at 100x g at 37°C. Cell sorting was performed on the Bio-Rad Sorter S3e instrument with the Software ProSort 1.5 (Bio-Rad, Hercules, California, USA).

### Molecular Cloning

Primers and plasmids used in this study are listed in **Table S10**. pMIG-FLAG-PTP1B-D181A (Medgyesi *et al*. 2014) was used as a PCR template to obtain pMIG-FLAG-PTP1B-D181A-Y46F. Two flanking PCR fragments containing the point mutation from tyrosine to phenylalanine and spanning two EcoRI restriction sites were amplified and joined by PCR. The fused product was ligated into the pJet1.2 cloning vector (Thermo Fisher Scientific, Waltham, MA USA). EcoRI (New England Biolabs) was used for restriction digest, to linearize the pMIG-FLAG-PTP1B-D181A vector and to release the point mutation containing insert from pJET1.2. Both were ligated to obtain pMIG-FLAG-PTP1B-D181A-Y46F.

pMIG-FLAG-PTP1B-Y46F Δ406-435 was cloned using pMIG-FLAG-PTP1B-D181A-Y46F as PCR template. The reverse primer was designed to introduce a stop codon and an EcoRI restriction site. The product was ligated into pJet1.2. followed by EcoRI restriction cloning as described above.

GST-tagged PTP1B for the peptide-based phosphatase assay was cloned using pMIG-FLAG-PTP1B. The vector was digested with BamHI and EcoRI and PTP1B was ligated into pGEX-2T (GE Healthcare, Pittsburgh, PA, USA).

The doxycycline inducible tet-on system comprises of two vectors: SH582 and SH461MK-EV-GFP (Haug *et al*. 2015), and was obtained from TET Systems GmbH & Co. KG, Heidelberg, Germany. The antibiotic resistance of the first vector was changed to bleomycin (SH582-bleomycin) by Gibson cloning (Gibson *et al*. 2009) using SH582 as backbone and pGIPZ (Open Biosystems, Huntsville, USA) for the bleomycin template. The second vector, pSH461MK-EV-GFP, was used as a control plasmid. Alternatively, the cDNA of FLAG-PTP1B, FLAG-PTP1B-D181A-Y46F, FLAG-PTP1B-D181A-Y46F-Δ406-435 was integrated into pSH461MK-EV-GFP using the corresponding pMIG-vector as a template for Gibson cloning. Carboxy-terminally Myc-tagged CD22 was generated by Gibson cloning. For this, pMSCV-TwinStrep_CD22 human (kind gift of AG Nitschke, Erlangen) (Meyer *et al*. 2020) and pMICD8 mb-1 FLAG (Infantino *et al*. 2010) were used as templates to amplify human CD22 and the pMICD8 backbone (Herzog and Jumaa 2007), respectively. The Myc-tag was introduced by the Gibson primers. The CD22 point mutations from tyrosine to phenylalanine were introduced by site-directed mutagenesis according to the QuikChange method (Agilent, Santa Clara, USA).

### CRISPR/Cas9-mediated PTP1B Gene Deficiency

To generate PTP1B KO cells, a targeted DNA double strand break was introduced using two guide RNAs, creating offset nicks with the Cas9n D10A nickase mutant of *Streptococcus pyogenes* (Le Cong *et al*. 2013). The vectors pX461 (Addgene) and C250 (derived from pX461, GFP replaced with dsRED (Lindner *et al*. 2017)) were digested with *Bbs*I (NEB) and ligated with the following annealed oligomers:

5’-CACCGTCCATCTCCATGACGGGCCA-3’ and 5’AAACTGGCCCGTCATGGAGATGGAC-3’ (first nick), 5’ CACCGATCGACAAGTCCGGGAGCT-3’ and 5’AAACAGCTCCCGGACTTGTCGATC

-3’ (second nick). Ramos B cells were electroporated to transfer 3 µg DNA of each purified plasmid [1350 V, 30 ms, 1 pulse, Neon Transfection System (Invitrogen)].

Expression of the gRNAs was monitored by flow cytometry (GFP and dsRED expression).

### Calcium measurements

For calcium measurements 1×10^6^ cells of each condition (Ramos cells treated with dimethylsulfoxid (DMSO) or with 0.5 µg/ml doxycycline for 2-3 days) were combined in a 1:1 ratio and loaded with 5 µg/ml Indo-1 (Molecular Probes, AAT Bioquest, Sunnyvale, USA) and 0.04% of pluronic F-127 (AAT Bioquest) in RPMI containing 1% FCS. Cells were incubated for 45 min at 37°C in the dark, washed twice with phosphate-buffered saline (PBS) and resuspended in 500 µl serum-free RPMI. Cells were incubated for 10 min at 37°C immediately before the calcium flux analysis on an LSR Fortessa instrument (Becton Dickinson, Franklin Lakes, USA) using the following parameters: Indo-1 “bound” 355 nm, BP 405/10 and “unbound” 355 nm, 445LP, BP 520/35; GFP 488 nm, BP 530/30. A baseline of 30 s was recorded before stimulation with 1 µg/ml anti-λ antibody (Southern Biotech, Birmingham, AL, USA). Flow cytometry standard files were analyzed using Flowjo^TM^ software for Windows, version 10.6.0 (Becton Dickinson) and its integrated kinetics tool. To quantify differences in the calcium flux curves of PTP1B KO *versus* GFP or FLAG-PTP1B a 5 point adjacent-averaging smoothing was performed using OriginPro 2017 (OriginLab Corporation, Northampton, MA, USA). An average of the first 30 s of the calcium measurement was calculated and used to normalize the measured values to the baseline. At timepoint 300 s the difference in the Indo-1 (bound/unbound) ratio was calculated between the curves of each sample. For the quantification of the calcium flux in the experiments with the different trapping mutants, the curves were baseline corrected as described above and the area under the curve of each sample was calculated.

### Immunofluorescence Microscopy

To study the subcellular localization of FLAG PTP1B WT, FLAG-PTP1B D181A Y46F or FLAG-PTP1B D181A-Y46F Δ406-435, doxycycline-inducible Ramos B cells were used. 2x 10^6^ cells were resuspended in 1 ml serum-free RPMI. 400,000 cells were seeded on each cover slip and fixed with ice-cold methanol/acetone (1:1, v/v) for 10 min at -20°C, followed by washing with 1x ice-cold PBS. Permeabilized cells were blocked with 10% normal goat serum and 1% bovine serum albumin (BSA) in PBS for 40 min at room temperature (RT). Subsequently, the cover slips were washed in 0.02% PBS-Tween 20 detergent (PBS-T), and cells were incubated with mouse anti-PTP1B FG6, Merck Millipore (1:100 in PBS-T) and rabbit anti-calnexin, Abcam (1:1,000 in PBS-T) for 2 h at RT. After washing (3x) with 0.02% PBS-T, samples were incubated with goat anti-mouse AlexaFluor 594 (1:200 in PBS-T) and goat anti-rabbit AlexaFluor 488 antibodies (1:200 in PBS-T) for 2 h at RT. Cell nuclei were stained with DAPI (1:1,000) for 2 min at RT. For mounting, ProLongGold (Life Technologies) was used. Images were obtained using a commercial gated STED microscope (TCS SP8 STED 3x) with a HC PL APO CS2 40x/1.30 oil objective (Leica Microsystems, Germany). Detection of DAPI, AlexaFluor 488 and 594 fluorophores was performed in sequential acquisition mode using PMT and HyD detectors in the standard mode and wavelengths ranges of 425-460, 503–550 and 601–649 nm, respectively.

### Proximity Ligation Assay

Proximity ligation assays (PLA) were performed as previously described (Kläsener *et al*. 2014). To generate PLA-probes against specific targets, the following unlabeled antibodies were used: anti-PTP1B (Merck Millipore MABS197, FG6-1G), anti-Igα (Novus Biologicals, HM47), anti-CD22 (Southern Biotech, RFB4) and anti-APLP2 (Proteintech Group, 15041-1-AP). The buffer of the anti-PTP1B antibody was exchanged to PBS-T using a Microcon-30kDa centrifugal filter unit (MRCF0R030, Merck Millipore) according to the manufacturer’s instructions. Anti-Igα Fab fragment was prepared with Pierce Fab Micro preparation kit (Thermo Scientific) using immobilized papain according to the manufacturer’s protocol. After desalting (Zeba spin desalting columns, Thermo Scientific), all antibodies were coupled with PLA Probemaker Plus or Minus oligonucleotides (Sigma-Aldrich) to generate PLA-probes. For *in situ* PLA, Ramos cells were settled on polytetrafluoroethylene slides (Thermo Fisher Scientific) for 30 min at 37 °C. Cells were fixed for 20 min with 4% paraformaldehyde, permeabilized with 0,5% saponin for 30 min at RT, and blocked for 30 min with blocking buffer (25 μg/ml sonicated salmon sperm DNA, 250 μg/ml BSA, 1M glycine). PLA was performed with Duolink *In Situ* Orange (Sigma Aldrich). Samples were directly mounted on slides with DAPI Fluoromount-G (SouthernBiotech) to visualize the PLA signals in relationship to the nuclei. Microscopic images were acquired with a Leica DMi8 microscope, 63 oil objective (Leica-microsystems). For each experiment a minimum of 1000 Ramos B cells from several images were analyzed with CellProfiler-3.0.0 (CellProfiler.org). Raw data were exported to Prism 7 (GraphPad, La Jolla, CA). For each sample, the mean PLA signal count per cell was calculated from the corresponding images and the statistical significance was determined using the Mann-Whitney test.

### Purification of Recombinant PTP1B

GST-PTP1B expression in BL21 competent E.coli (NEB) was induced by the addition of 0.2 mM IPTG. After incubation at 37°C for 3 h, bacteria were pelleted at 7000 g for 10 min. Cell lysis was performed for 1 h on ice in PBS supplemented with 10 mM MgCl_2_, 1 mM PMSF, 0.1 mg/ml lysozyme and 5 U/ml benzonase and completed by 10 min sonification on ice. The lysate was cleared by centrifugation for 40 min at 25,000 g at 4°C. GST-PTP1B was purified with a 1 ml GSTrap FF Column on an ÄKTA pure purification system (GE Healthcare GmbH.) using a binary buffer system (buffer A: 50 mM HEPES, pH 8.5, 150 mM NaCl; buffer B: 50 mM HEPES, pH 8.0, 10 mM reduced glutathione). The lysate was applied at a flowrate of 1 ml/min. A linear gradient from 0-100% buffer B in 10 column volumes was applied to elute the sample. The eluate was collected in 0.5 ml fractions. PTP1B-containing fractions were pooled and supplemented with 20% glycerol.

### Materials for Peptide Synthesis and Purification

All amino acids and resins were purchased from Novabiochem (Merck, Darmstadt, Germany) and Carbosynth (Compton, United Kingdom). All other synthetic reagents were obtained from Novabiochem, Sigma-Aldrich (St. Louis, Missouri, USA), or Carl Roth (Karlsruhe, Germany). Peptide synthesis was performed on a MultiPep RSi peptide synthesizer (Intavis Bioanalytical Instruments AG, Köln, Germany). Peptides were purified using a 1260 Infinity II preparative HPLC system (Agilent Technologies, Santa Clara, California, USA) with a VP 250/10 NUCLEODUR 100-5 C18 ec column (Macherey-Nagel, Düren, Germany) running a general gradient of 10% to 70% acetonitrile (ACN) in H_2_O and validated using a 1260 Infinity I HPLC System coupled to a 6120 Quadrupole LC/MS (Agilent Technologies, Santa Clara, California, USA) with an EC 250/4 NUCLEODUR 100-5 C18 ec column (Macherey-Nagel) with a general gradient of 10% to 90% ACN in H_2_O, and additionally a Microflex LT MALDI (Bruker, Bremen, Germany) was used for the analysis.

### Peptide Synthesis

Peptides were synthesized following fluorenylmethyloxycarbonyl-(Fmoc) solid-phase peptide synthesis (SPPS) (**Table S10**). Pre-coupled, tert-butyloxycarbonyl protected chlorotrityl resins were used for all peptides. Resins were swollen for 20 min in N,N-dimethylformamide (DMF) prior to synthesis in the peptide synthesizer. One round of synthesis was carried out for each amino acid in the peptide sequence. A synthetic round consisted of double coupling to an Fmoc and side chain-protected amino acid residue, followed by a capping step with acetic anhydride (Ac_2_O), and then removal of the Fmoc group with piperidine in preparation for the next round of coupling. Coupling reactions were carried out by adding Fmoc-protected amino acids (4 eq.), O-benzotriazole-N,N,N’,N’-tetramethyl-uronium hexafluorophosphate (HBTU) (4 eq.), Nhydroxybenzotriazolehydrate (HOBt) (4 eq), and N-methylmorpholine (NMM) to the resin in 1 ml DMF and reacting for 30 - 45 min. Capping was done by reaction with a 1 ml solution of 5% Ac_2_O and 5% 2,6-Lutidine in DMF for 5 min. Fmoc-deprotection was achieved by addition of a 20% piperidine in DMF solution to the resin, first for 3 min, then again for 8 min. Resins were washed with DMF between each step. After Fmoc removal of the final amino acid residue to reveal the free N-terminus, peptides and libraries were removed from the resin and fully deprotected in one step by shaking overnight in cleavage cocktail (95% trifluoroacetic acid (TFA), 2.5% triisopropylsilane (TIPS), and 2.5% H_2_O). Peptides were precipitated in cold ether (-20°C) and collected by centrifugation. Peptides were then validated by MALDI and HPLC-MS, and purified by preparative HPLC.

### Detection of Phosphate Release from Peptides

The peptides were dissolved in 25% DMSO and 75% H_2_O at 7.5 mM after purification. The EnzChek® Phosphate Assay Kit (Thermo Fisher Scientific) was used to assess the kinetics/dynamics of the peptide dephosphorylation by recombinant PTP1B. The assay was carried out in a volume of 100 μL reaction buffer (20 mM Tris, 100 mM NaCl, 2 mM DTT, 0.15 units purine nucleoside phosphorylase, 0.2 mM 2-amino-6-mercapto-7- methylpurine riboside) containing indicated concentrations of peptide and 20 nM recombinant PTP1B. The change of absorbance was detected at 360 nm and 28°C on a Synergy H1 microplate reader. For analysis of enzyme kinetics, raw data was further analyzed in GraphPad Prism v6. Error bars represent SD of three replicates. Values were compared to a standard curve prepared by using the phosphate-solution included in the kit and following the manufacturer’s instructions.

### B Cell Stimulation and Cell Lysis

For stimulation of Ramos B cells, 1×10^7^ cells were starved for 30 min at 37°C in 1 ml FCS-free RPMI and then treated with 10 µg/ml anti-λ antibody (Southern Biotech) under shaking at 750 rpm for the indicated time points. To stop the reaction, cells were spun down for 10 s at 21,000xg, the supernatant was removed and the cell pellet was immediately frozen in liquid nitrogen. Cell lysis was performed with modified RIPA buffer (1x PBS, 1% Nonidet P-40, 0.1% SDS, 0.5% sodium deoxycholate), supplemented with 2 mM sodium orthovanadate, 10 mM sodium pyrophosphate, 9.5 mM sodium fluoride, 10 mM ß-glycerophosphate and 1x EDTA-free protease inhibitor cocktail (Roche) prior to lysis.

### Co-Immunoprecipitation

2×10^8^ cells were lysed for 25 min in 5 ml ice-cold lysis buffer (50 mM Tris-HCl, pH 8.5, 137.5 mM NaCl, 1 mM EDTA, 1% Triton X-100, 1 mM sodium fluoride, EDTA-free protease inhibitor cocktail [Roche]). 1 mM orthovanadate was added to the lysis buffer to prevent binding of substrates to the PTP1B trapping mutant. Lysates were cleared by centrifugation for 10 min at 4,969xg. A small aliquot was taken for Western blot analysis of the lysate, the remaining sample was mixed with 40 µl FLAG M2 magnetic beads (Sigma). The samples were incubated for 2-4 h on a rotating wheel (10 rpm) at 8°C, washed 3x with 1 ml lysis buffer and eluted with 40 µl 1x sample buffer (10% glycerol, 1% β-mercaptoethanol, 2% SDS, 50 mM Tris-HCl, pH 6.8, 0.004% bromophenol blue). For MS analysis, the proteins were eluted with 40 µl 3x FLAG peptide (Sigma-Aldrich) and collected by centrifugation for 30 min at 100xg.

For the phosphatase assay, Ramos B cells stably expressing CD22-Myc were used. Expression of FLAG-PTP1B was induced 18 to 24 h prior to the assay by the addition of doxycycline as indicated. Cells were stimulated with 10 µg/ml anti-λ for 3 min at 37°C or left serum starved and lysed in 50 mM Tris-HCl pH 8.0, 1% Triton X-100, 137.5 mM NaCl, supplemented with 2 mM sodium orthovanadate, 10 mM sodium pyrophosphate, 1 mM sodium fluoride, 10 mM ß-glycerolphosphate and 1x protease inhibitor cocktail (Roche). For immunoprecipitation, Pierce™ anti-c-Myc magnetic beads (Thermo Fisher Scientific) were used.

### In-Gel Protein Digestion

Co-IP samples were loaded and separated on a 4-20% Tris-Glycine gel (Invitrogen). Proteins were stained with colloidal Coomassie Brilliant Blue G-250. In gel-digestion of proteins was performed essentially as described before (Schwarz *et al*. 2015) with the exception that protein thiols were reduced in 5 mM tris (2-chloroethyl)phosphine (TCEP) for 10 min at 60°C and alkylated in 100 mM 2-chloroacetamide (CAM) for 15 min at 37°C.

### Western Blot Analysis

FLAG-tagged PTP1B was detected with anti-FLAG antibody (20543-1-AP (Proteintech, Manchester, UK)). PTP1B was detected with anti-PTP1B clone FG6-1G (MABS197, Merck Millipore) or with clone H-135 (sc-14021) from Santa Cruz Biotechnology to detect specifically the C-terminal part. Anti-CD22 antibody was purchased from Novus Biological (clone 2H1C4), anti-CD79A (Igα; clone HM47) and anti-SYK (4D10.2) from Biolegend. The antibodies directed against; BTK-pY223, #5082; PLCγ1, #5690; PLCγ2, #3872; PLCγ2-pY759, #3874; SYK-p525/526, #2710; ERK, #4695; ERK-pT202/Y204, #4370; GRB2, #3972; and GAPDH, #2118 were all from Cell Signaling. Anti-phospho- CD22-pY807 (ab32040) antibody was purchased from Abcam. Anti-Myc-tag (66004-1-Ig), anti-APLP2 (15041-1-AP) and anti-CBL (25818-1-AP) were from Proteintech. Anti-BTK (sc-1107) and anti-BCAP (AF4857) antibodies were obtained from Santa Cruz Biotechnology and R&D Systems (Inc., Minneapolis, USA), respectively. All antibodies were used according to the manufacturer’s instructions. For imaging, a ChemoCam (Intas, Göttingen, Germany) equipped with a full-frame 3.2 megapixel Kodak KAF-3200ME camera was used. Western blot signals were quantified with Quantity One 4.6.9 (Bio-Rad, Hercules, CA). For statistical analysis, two sample t-tests were performed over three replicates using Microsoft Excel 2013. Error bars represent the standard error of the mean (SEM).

### Sample Preparation for Quantitative Proteome Analysis

Ramos B cells were harvested and washed three times with ice-cold 1x PBS. Cells were lysed in 30 mM Tris base with 7 M urea and 2 M thiourea, pH 8.5, supplemented with protease inhibitor (Roche). To aid cell lyses and to sheer DNA, cells were sonicated on ice at least twice with an ultrasonic homogenizer for 10 s at 90% of maximum power. Cell debris was removed by centrifugation for 20 min at 21.000 x g and 4°C. Samples containing 300 µg of total protein were prepared and cysteine residues were reduced with 5 mM TCEP for 20 min at room temperature (RT) and alkylated with 50 mM CAM for 20 min at 37°C. The reaction was quenched with 10 mM dithiothreitol (DTT). Each sample was then diluted 1:1 with 50 mM ammonium bicarbonate and digested with Lys-C (protein to enzyme ratio 1:100, FUJIFILM Wako Chemicals Europe GmbH, Neuss, Germany) for 4 h at 37°C Subsequently, the sample was diluted 1:1 with 50 mM ammonium bicarbonate and proteins were digested with trypsin (protein to enzyme ratio 1:50, Promega, Mannheim, Germany) overnight at 37°C. The reaction was stopped by acidification with TFA (1% [v/v] final concentration) and the peptide sample was desalted using a Empore^TM^ Solid Phase Extraction cartridge (3M Bioanalytical Technologies, St. Paul, USA). The cartridge was washed with 1 ml methanol followed by 1 ml 70% ACN with 0.1% TFA. The column was conditioned twice with 0.5 ml 0.1% TFA before the sample was applied. Peptides were washed with 0.5 ml 0.1% TFA and eluted with 0.5 ml 70% ACN. Peptides were freeze-dried overnight using an Alpha 1-2 LDplus system (Christ, Osterode am Harz, Germany)

### High-pH Reversed-phase Fractionation

High-pH reversed-phase fractionation was performed on an Ultimate 3000 HPLC system equipped with a Gemini-NX 3µ C18 110 Å column (Phenomenex, Aschaffenburg, Germany). The desalted sample was taken up in 300 µl of 10 mM NH_4_OH (solvent A). A constant flow rate of 0.2 ml/min was applied and peptides were loaded onto the column at 1% solvent B (10 mM NH_4_OH in 90% ACN) for 5 min. Peptides were separated applying a linear gradient from 1% B to 61% B in 55 min, followed by an increase to 78% B in 2 min, a decrease to 1% B in 5 min, and re-equilibration of the column for 16 min. Fractions were collected every 43 s, starting at 1.5 min until 70 min. Of the collected 96 fractions, every 32^nd^ fraction was pooled.

### High-Performance Liquid Chromatography and Mass Spectrometry

Reversed-phase HPLC was performed on an UltimateTM 3000 RSLCnano system (Thermo Fisher Scientific, Dreieich, Germany) equipped with two PepMap^TM^ C18 μ-precolumn (0.3 mm × 5 mm, 5 µm, 300 Å Thermo Fisher Scientific) and an Acclaim^TM^ PepMap^TM^ analytical column (75 μm × 250 mm, 3 μm, 100 Å, Thermo Fisher Scientific). For LC-MS analysis of affinity-purified samples (SILAC Co-IP), the RSLC system was coupled to a Velos Orbitrap Elite instrument (Thermo Fisher Scientific, Bremen, Germany). A binary solvent system was used with solvent A consisting of 0.1% formic acid (FA) and 4% DMSO and solvent B consisting of 48% methanol, 30% ACN, 0.1% FA and 4% DMSO. Peptides from SILAC Co-IPs were washed and concentrated for 5 min with 0.1% TFA on the pre-column. The gradient applied for peptide separation on the analytical column (flow rate of 0.250 µl/min) was as follows: 1% to 65% solvent B in 50 min, increased to 95% B in 5 min, 5 min at 95% B, followed by a decrease to 1% B in 1 min. Subsequently, the column was re-equilibrated for 14 min with 1% B. The Velos Orbitrap Elite instrument was operated with the following parameter settings: spray voltage, 1.5 kV; capillary voltage, 200 V; automatic gain control (AGC), 1×10^6^ ions; max. ion time (IT), 200 ms. Full scans were acquired in the orbitrap with a mass range of *m*/*z* 370 to 1,700 and a resolution (R) of 60,000 at *m/z* 400. A TOP15 method was applied for collision-induced dissociation (CID) of multiply charged peptides performed in the linear ion trap with a normalized collision energy (NCE) of 35% and an activation q of 0.25. The AGC for CID was set to 5,000, the max. IT was 150 ms and the dynamic exclusion time (DE) for previously selected precursor ions was 45 s.

For quantitative proteome analysis, the RSLC system was coupled to a Q Exactive Plus instrument (Thermo Scientific). A solvent system consisting of 0.1% FA (solvent A) and 86% ACN, 0.1% FA (solvent B) was used for peptide separation. The RSLC was operated with a flow rate of 0.250 µl/min for the analytical column. The gradient was 4% to 40% B in 50 min, followed by 40 to 95% B in 5 min. After 5 min at 95%, the column was re-equilibrated for 15 min with 4% B. Full scans were acquired in the orbitrap in a mass range of *m*/*z* 375 to 1,700 with the following parameters: R, 70,000 at *m/z* 200; AGC, 3×10^6^ ions; max. IT, 60 ms. For MS/MS experiments, a TOP12 method was used and parameters were as follows: NCE, 28%; DE 45 s; AGC, 5,000; max. IT, 120 ms.

### Bioinformatics

Raw data were searched against the Uniprot Human Proteome set (release 2017_06, 93,591 protein entries) using MaxQuant 1.5.5.1 (for SILAC Co-IP and for global proteome analysis in Ramos B cells) or version 1.5.4.0 (quantitative proteome analysis of PTP1B KO Ramos B cells) with its integrated search engine Andromeda (Cox and Mann 2008; Cox *et al*. 2011). The species was restricted to *homo sapiens* as Ramos B cells are of human origin. MaxQuant default settings were used unless stated otherwise. Database searches were conducted with trypsin (for SILAC Co-IPs) or with trypsin and Lys-C (for quantitative proteome analysis) as proteolytic enzymes, a maximum number of three missed cleavages, and mass tolerance of 4.5 ppm for precursor and 0.5 Da (CID data) or 20 ppm (HCD data) for fragment ions. For the quantitative proteome analysis of PTP1B KO cells, two missed cleavages were allowed. Carbamidomethylation of cysteine residues was set as fixed modification, and oxidation of methionine, N-terminal acetylation were applied as variable modifications and additionally serine/threonine/tyrosine phosphorylation for SILAC Co-IPs and for global proteome analysis of Ramos B cells. The minimum number of unique peptides was set to 1. A false discovery rate (FDR) of 1% was applied to both peptide and protein lists. For global proteome analysis, “label-free” and “iBAQ” quantification as well as “match between runs” were enabled with default settings. For SILAC Co-IPs, the multiplicity was set to 3 and “Requantify” was enabled. The maximum number of labelled amino acids was set to 4. For the quantitative proteome analysis of PTP1B KO cells, multiplicity was set to 2, the maximum number of labelled amino acids to 3 and “Requantify” was enabled. For protein quantification, “razer and unique peptides” were used with a minimum ratio count of 2. For the determination of significantly enriched PTP1B substrate candidates, non-normalized SILAC ratios were log_10_-transformed and a mean log_10_ ratio over at least 2 out of 3 replicates was calculated. P-values were determined with a right-sided t-test in Perseus 1.5.5.3 (Tyanova *et al*. 2016). For Gene Ontology (GO) and REACTOME pathway enrichment analysis, the Cytoscape 3.2.1 (Shannon *et al*. 2003) plugin ClueGO v2.5.1 (Bindea *et al*. 2009) was used. The global proteome dataset was divided into five quantiles with an equal number of protein groups according to their abundance rank. Enrichment of each quantile against the human reference set was analysed for the GO domain “biological process” (BP) and for REACTOME pathways. A right-sided hypergeometric statistic test with Bonferroni step down FDR correction at a significance level of 0.05 was used for p value calculation.

For network analysis of PTP1B substrate candidates identified by SILAC Co-IPs, the STRING application in Cytoscape 3.8.2 was used. For network analysis of the Ramos B cell proteome, the pathway was downloaded from NetPath (Kandasamy *et al*. 2010) and processed in Cytoscape 3. 2. 1. Enrichment of SH3 domain containing proteins was calculated using the SMART database (Letunic *et al*. 2015; Letunic and Bork 2018). For the quantitative proteome analysis of PTP1B KO cells, normalized SILAC ratios were log_2_-transformed, a mean log_2_ ratio was calculated over three out of four biological replicates and a two-sided t-test was applied to determine p-values. Extracted ion chromatograms of PTP1B peptide precursor ions were analyzed manually in Skyline 3.5.0 (Schilling *et al*. 2012).

### Protein Copy Number Determination

To determine protein copy numbers, the proteomic ruler plugin v.0.1.6 (available from http://www.coxdocs.org/doku.php?id=perseus:user:plugins:store) for Perseus 1.5.5.3 was used (Tyanova *et al*. 2016). The applied settings were as follows: averaging mode, “same normalization for all columns”; detectability correction, number of detectable tryptic peptides; quantification, histones as proteomic ruler; ploidy of the cells, 1.9565 as Ramos B cells have 45 chromosomes. From the resulting copy numbers of three biological replicates, mean log_10_ copy numbers were calculated for protein groups with at least two valid values.

### RNA Sequencing Data Analysis

RNA sequencing raw files from Qian and colleagues (Qian *et al*. 2014) were downloaded from the sequence read archive (SRA) with the following accession numbers: SRX753158 (replicate 1), SRX753159 (replicate 2) and SRX753160 (replicate 3). The quality of the raw data was assessed with FastQC v0.11.5 (https://www.bioinformatics.babraham.ac.uk/projects/fastqc/). Raw reads had a length of 50 bases and were trimmed with BBDuk v.37.28 implemented into Geneious v11.0.5 [http://www.geneious.com, (Kearse *et al*. 2012)] on both ends, if the quality of the base call was less than 20 phred score. Reads shorter than 45 bases were discarded. Remaining reads were mapped against Homo sapiens GRCh38.p7 primary assembly with the Geneious RNA assembler using default settings at medium sensitivity and with the minimum mapping quality adjusted to 99.9% confidence. Expression levels (RPKM and TPM) were calculated for each replicate with Geneious v11.0.5. As a measure for transcript abundance, transcript per million (TPM) was used (Wagner *et al*. 2012). TPM values for each replicate were log_2_-transformed and mean log_2_ TPM values were calculated. Resulting data were binned into a histogram with Origin 2017 (OriginLab, Northampton, MA, USA). A cut-off value of log_2_ TPM = 0 was set for genes considered to be expressed. Entries with non-zero TPM values were mapped against the Uniprot database according to their GeneID. Non-mapping entries were further mapped according to their gene name to Uniprot identifiers. All mapped entries were considered as protein-coding genes.

### Data availability

All raw data and original MaxQuant result files have been deposited to the ProteomeXchange Consortium (http://proteomecentral.proteomexchange.org) via the PRIDE partner repository (Perez-Riverol et al., 2018) with the dataset identifiers PXD024037 (large-scale quantitative analysis of human Ramos B cells), PXD024086 (quantitative PTP1B substrate trapping) and PXD024038 (quantitative analysis of Ramos B cells *versus* Ramos B cells with PTP1B KO).

## ACKNOWLEDGEMENTS

We thank the PRIDE team for data deposition to the ProteomeXchange Consortium, Bettina Knapp for technical assistance in LC-MS analysis and Lars Nitschke and Sarah Meyer for scientific discussions. Work included in this study has been performed in partial fulfilment of the requirements for the doctoral thesis of J.J.S., and the master thesis of S.F. and L.G. at the University of Freiburg. This work was supported by the Deutsche Forschungsgemeinschaft (DFG, German Research Foundation) TRR 130 project P05 (to B.W. and M.R.) and project C02 (to B.W.), Project ID 403222702/SFB 1381 (to B.W. and M.K.), FOR 2743 (to B.W. and M.K.) as well as Germany’s Excellence Strategy (CIBSS – EXC-2189 – Project ID 390939984; to B.W., M.R. and M.K.) and the Excellence Initiative of the German federal and state governments (EXC 294, BIOSS; GSC-4, Spemann Graduate School; to B.W. M.R. and M.K.)

## AUTHOR CONTRIBUTION

J.J.S. performed proteomic, bioinformatic, protein interaction and biochemical analyses with the support of S.F. and D.M. L.G. performed the PTP1B phosphatase assay. T.K. the peptide-based phosphatase assay and K.K. the PLA study. All authors analyzed data, designed and interpreted experiments. B.W. supervised the study. B.W. and M.R. conceived the project. B.W. and J.J.S. wrote the manuscript with the input of M.R. All authors approved the final version of the manuscript.

## SUPPLEMENTAL INFORMATION

**Supplemental Figure 1.**
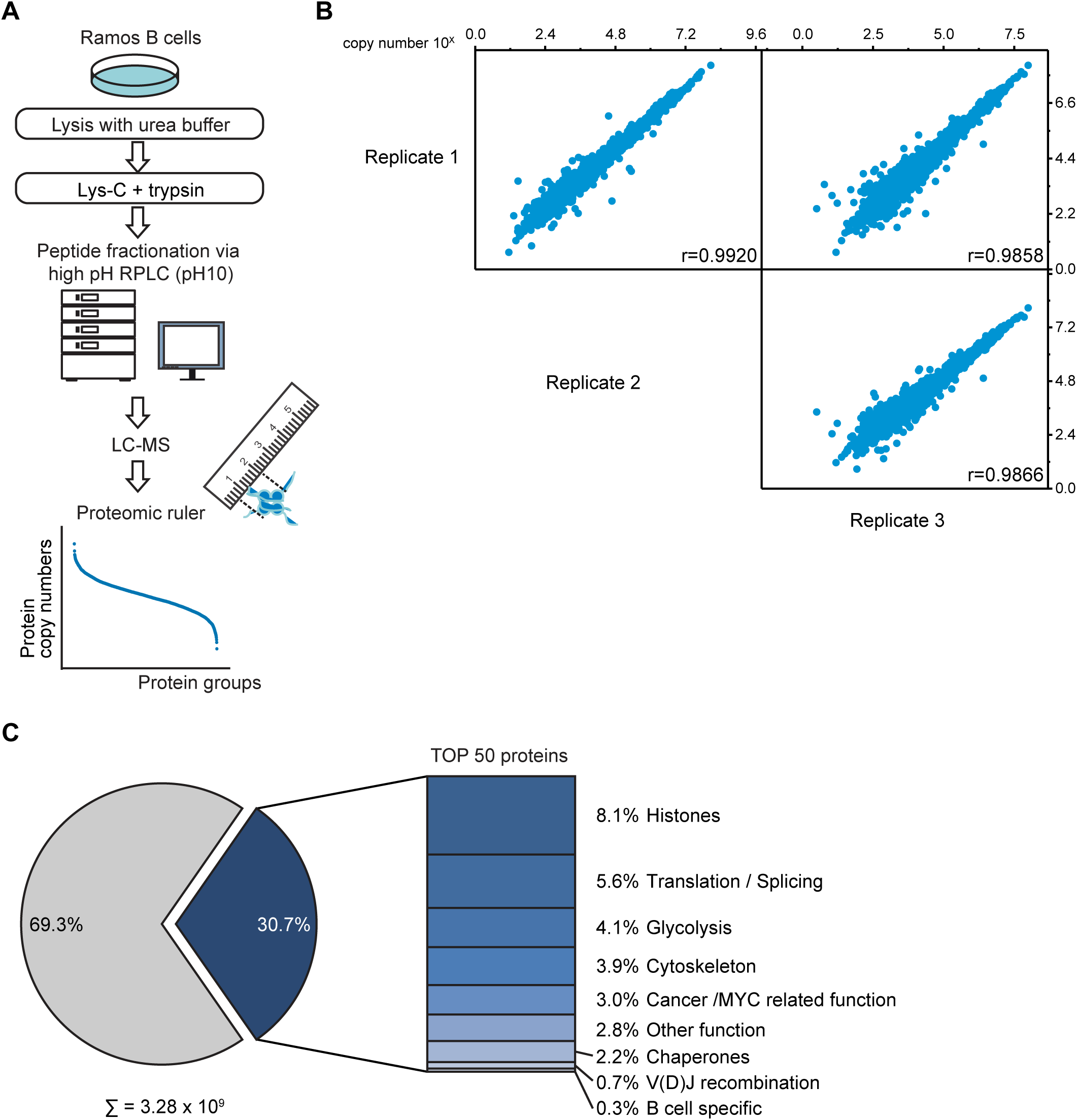
Absolute quantitative analysis of the Ramos B cell proteome. Related to Figure 1 **A.** Workflow of the global absolute quantitative proteome analysis. Ramos B cells were lysed in urea buffer and proteins digested in-solution using Lys-C and trypsin. Proteolytic digests were fractionated using high-pH reversed-phase liquid chromatography (RPLC) and analyzed by LC-MS. Copy numbers of proteins were calculated based on MS1 data from three independent biological replicates using the Proteomic ruler method (Wiśniewski *et al*. 2014). **B.** Multi-scatterplot depicting the correlation of the determined log_10_ copy number of proteins per Ramos B cell between individual replicates. Pearson’s correlation coefficient, r. **C.** Piechart depicting the share of the 50 most abundant proteins on the total copy number per Ramos B cell. The bar chart shows the classification of these 50 proteins in different functional categories (see Table S1).

**Supplemental Figure 2.**
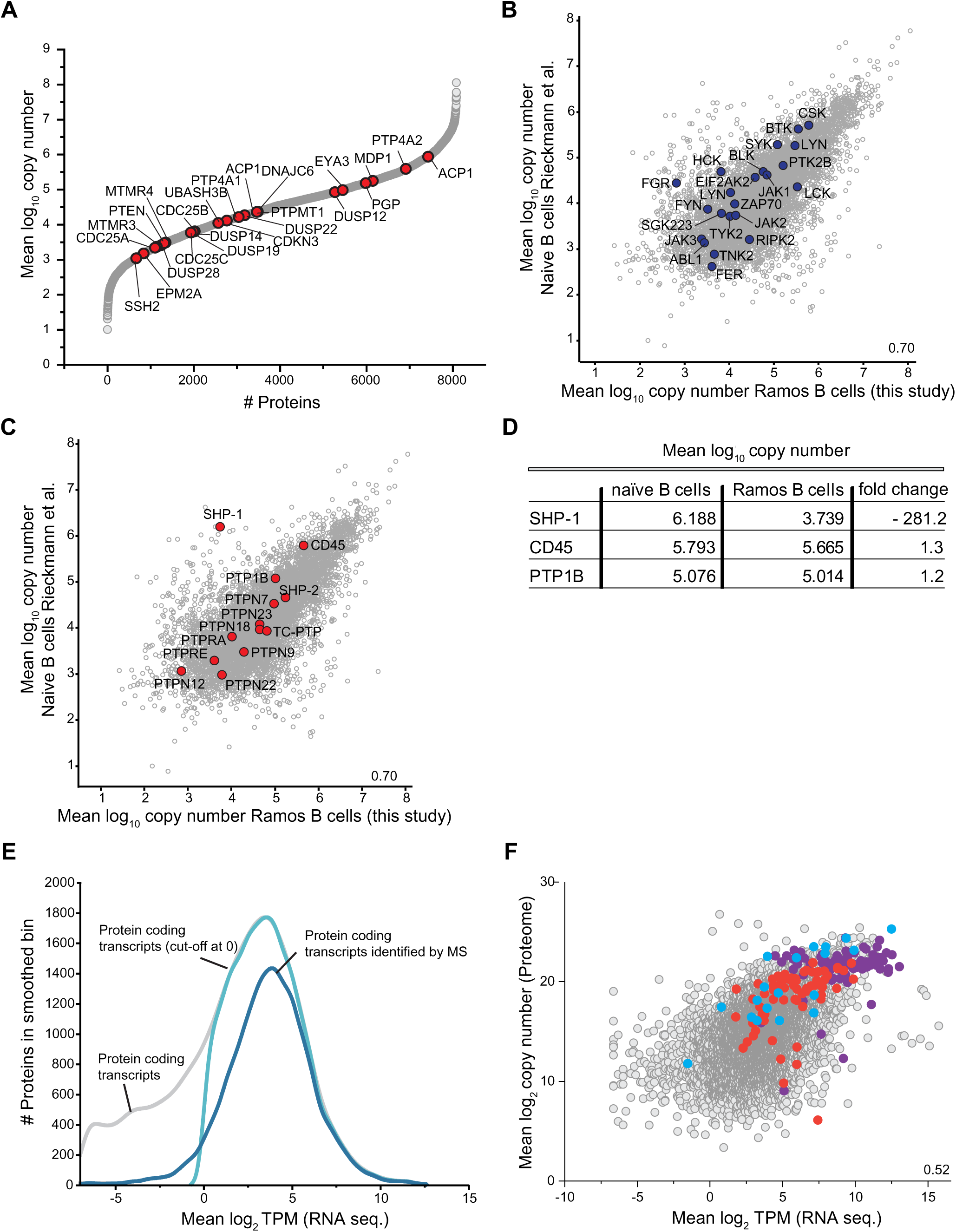
Evaluation of absolute quantitative proteome data of human Ramos B cells. Related to Figure 2 **A.** Protein copy number profile of protein tyrosine phosphatases (PTPs), without the “classical” PTPs shown in Fig. 2B, identified in Ramos B cells. **B, C.** Scatterplots comparing mean log_10_ protein copy numbers of naïve B cells ((Rieckmann *et al*. 2017)) and Ramos B cells reported in this study. Protein tyrosine kinases present in the dataset are depicted in blue (B) and “classical” PTPs in red (C). **D.** Mean log_10_ copy numbers of the three most abundant “classical” PTPs in naïve B cells compared to Ramos B cells and the difference in expression given as fold-change. **E.** Kernel-smoothed distribution curves of RNA sequencing (seq.) data of human Ramos B cells calculated from raw data reported in (Qian *et al*. 2014). Grey, distribution of transcripts of protein-coding genes; light blue, distribution of transcripts of protein-coding genes with a log_2_ mean transcript per million (TPM) value > 0 (cut-off value of 1); dark blue, protein-coding transcripts for which proteins were identified by MS analysis in this work (see Figure 1 and S1 and Table S1). **F.** Comparison of log_2_-transformed mean TPM values and protein copy numbers of ribosomal proteins (purple), proteasome components (red) and enzymes of canonical glycolysis (blue).

**Supplemental Figure 3.**
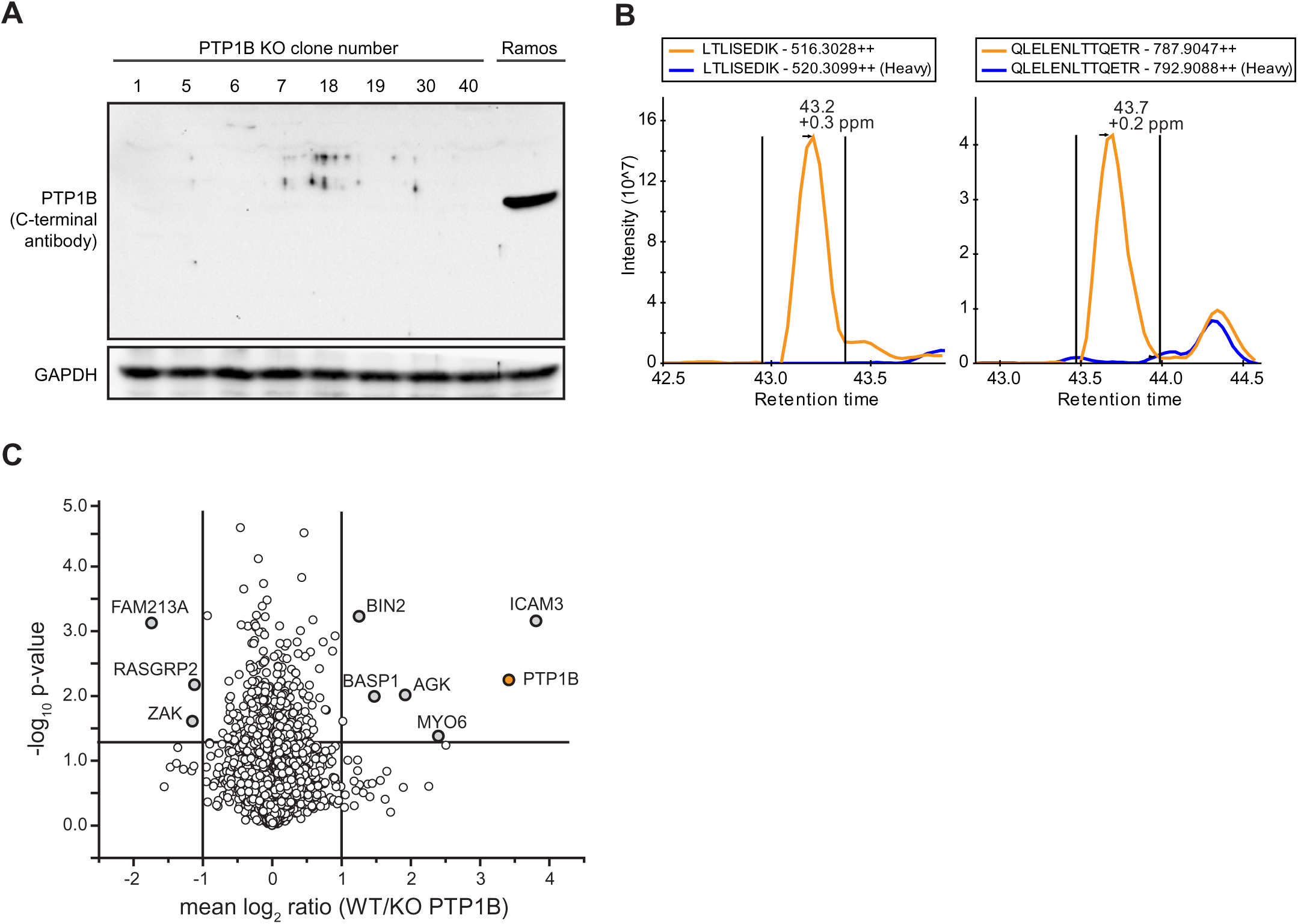
Verification of PTP1B knockout in human Ramos B cells. Related to Figure 3 **A.** Western blot analysis of different PTP1B knockout (KO) clones (clone numbers are indicated). The loss of PTP1B protein was controlled with an antibody directed against the C-terminal part of PTP1B. As control, non-gene-edited Ramos B cells (Ramos) were used. Equal loading of samples was monitored by the detection of GAPDH. **B.** Extracted ion chromatograms (XICs) of selected doubly charged unique peptides of PTP1B. Indicated in orange and blue are the XICs of the light- and heavy-labeled peptide versions originating from non-gene-edited and gene-edited PTP1B KO Ramos B cells. The data confirm the absence of PTP1B at the protein level in the KO cells. **C.** Quantitative MS analysis to confirm the loss of PTP1B at the protein level in PTP1B knockout (KO) Ramos B cell lines. Ramos B cells and two PTP1B KO cell lines made from two different clones (clone 1 and 5) were differentially labeled with stable isotope-coded amino acids (SILAC technology) and proteins digested in solution-digestion using trypsin and LysC followed by high pH-reversed phase fractionation and quantitative LC-MS analysis. Mean log_2_ ratios of Ramos B cells and PTP1B KO cells were calculated over three out of four biological replicates and a two-sided t-test was applied to determine p-values. PTP1B is indicated in blue and proteins with a minimum fold-change of 2 and a p-value ˂ 0.05 were labeled. For PTP1B, a fold-change of 3.42 was calculated by MaxQuant (Cox and Mann 2008).

**Supplemental Figure 4.**
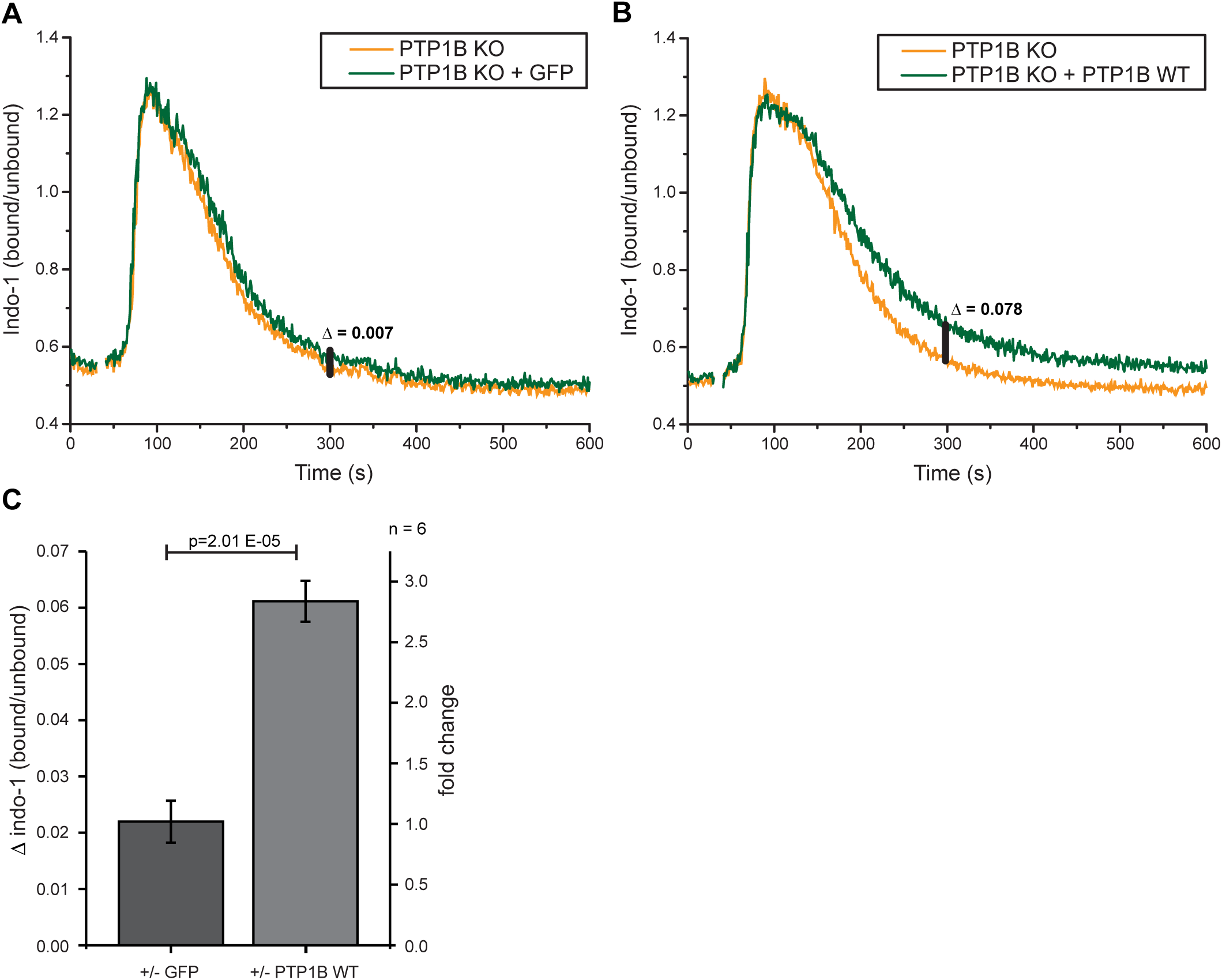
PTP1B re-introduction delays the return of the intracellular calcium level to the baseline. Related to Figure 3 **A, B.** Intracellular calcium measurements were performed using PTP1B knockout (KO) cells with inducible expression of either empty vector control (GFP only) or FLAG-PTP1B wildtype (WT). Basal calcium levels were recorded for 30s before the addition of 1 µg/ml anti-λ to stimulate BCR-induced calcium mobilization. Δ indicates the baseline corrected difference of the indo-1 (bound/unbound) ratio between the samples at 300s. **C.** Quantification of the calcium response in B cells at 300s. Statistical analysis was performed using a student’s t-test (two-sided, n=6, error bars = SEM).

**Supplemental Figure 5.**
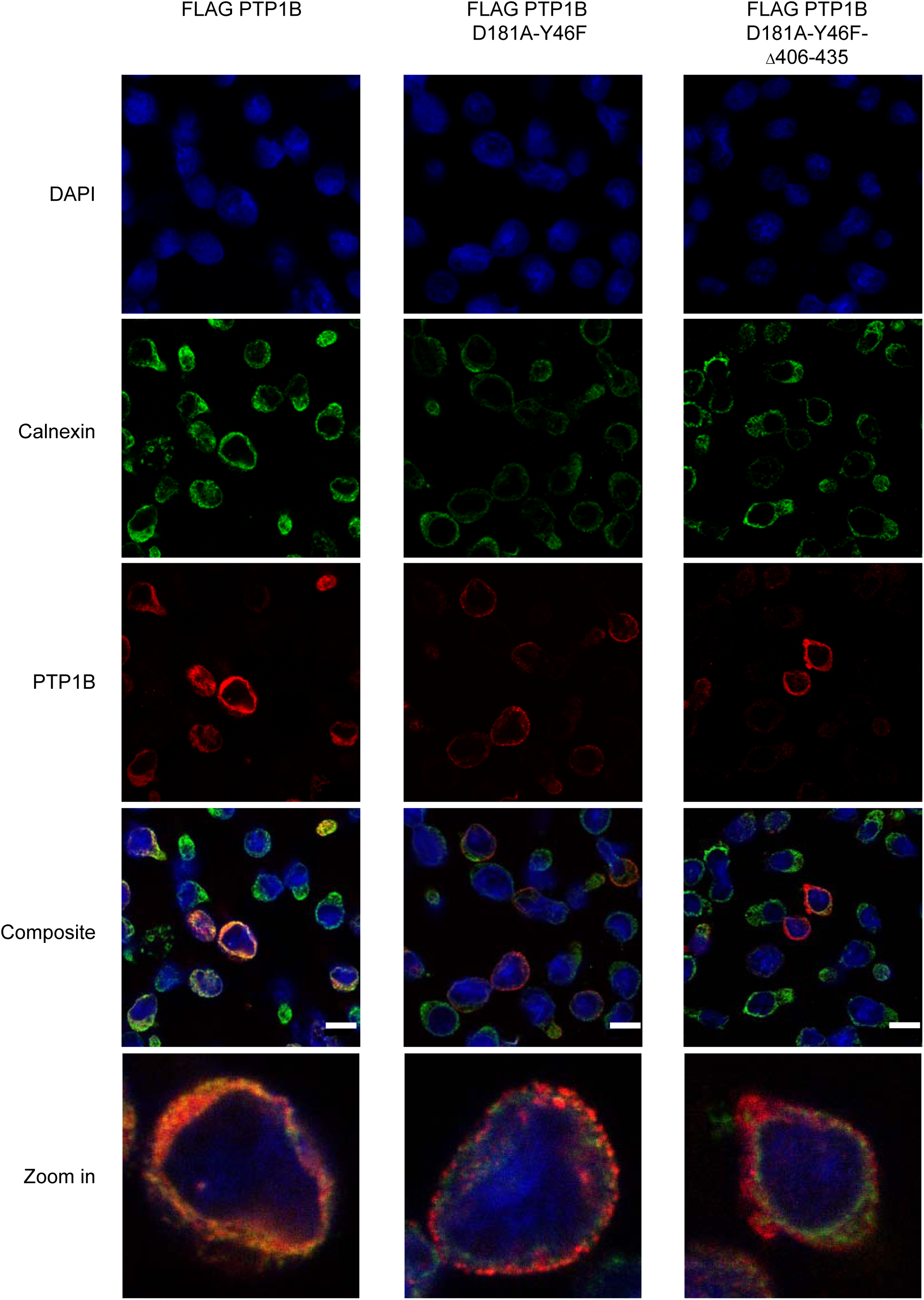
Analysis of FLAG-tagged PTP1B and PTP1B substrate mutant versions in human Ramos B cells by immunofluorescence microscopy. Related to Figure 4 **A.** Localization analysis of FLAG-PTP1B, FLAG-PTP1B-D181A-Y46F and FLAG-PTP1B-D181A-Y46F d406-435 in Ramos PTP1B KO cells by immunofluorescence microscopy. The expression of the constructs was induced by doxycycline 24 h prior to the experiment. Cells were seeded on cover slips, fixed and stained with DAPI (nucleus, blue), anti-calnexin antibody (ER marker, green) and anti-PTP1B antibody (PTP1B, red). Shown are representative excerpts of acquired images. For better visualization, single cells were zoomed in. FLAG-PTP1B WT localized mainly to the ER, as shown by overlay with the ER marker protein calnexin. The FLAG-PTP1B trapping mutant only marginally overlapped with the ER marker with a punctated ring-like structure. The ER domain-lacking PTP1B trapping mutant did not overlap with the ER marker and showed a diffuse staining, pointing to its expected cytosolic location in Ramos B cells. Scale bar, 10 µm

**Supplemental Figure 6.**
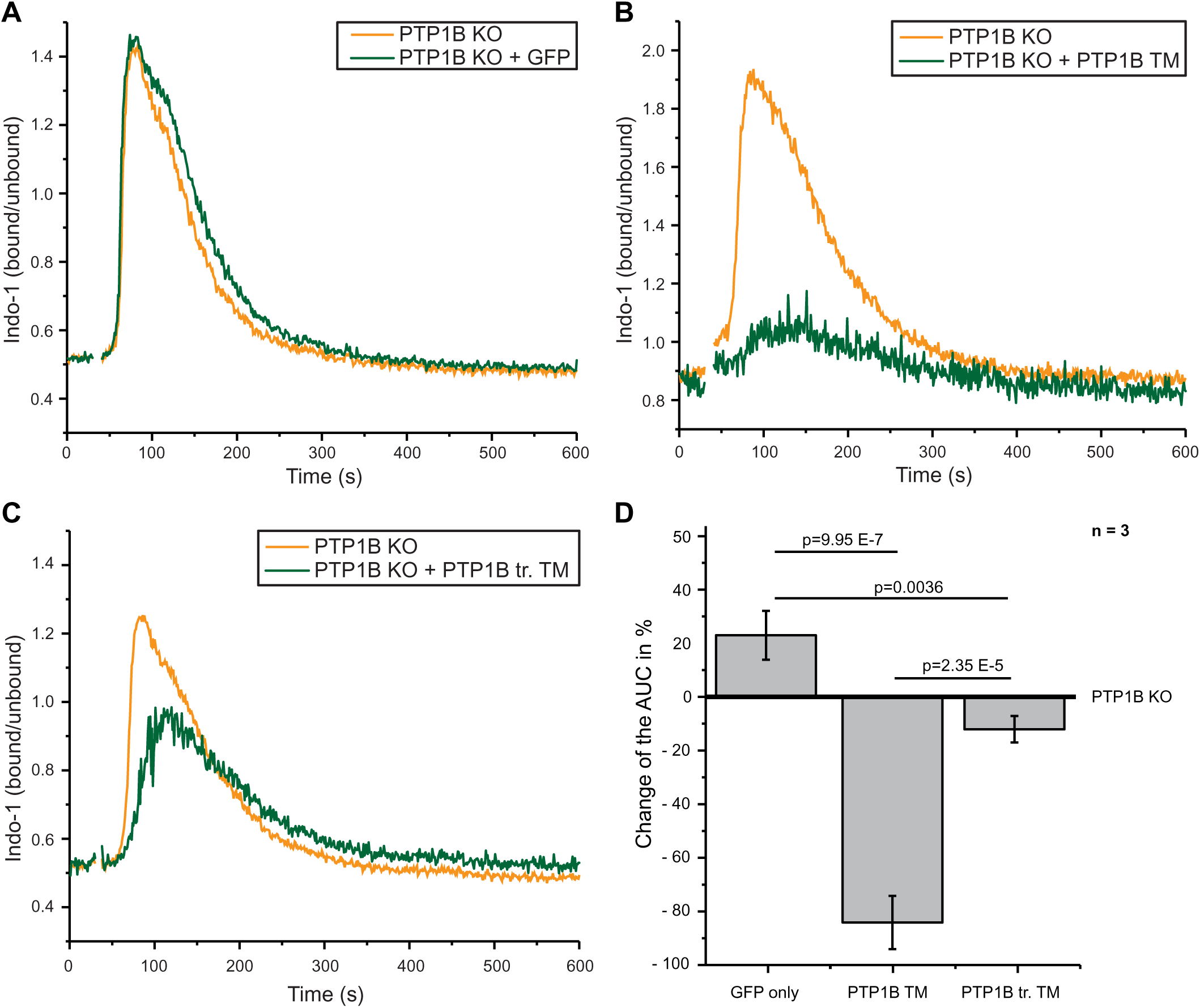
PTP1B trapping mutants capture proteins involved in Calcium signalling. Related to Figure 5 **A-C.** Intracellular calcium mobilization after anti-λ stimulation (1 µg/ml) in PTP1B knockout cells with inducible expression of empty vector control (GFP only), FLAG-PTP1B-D181A-Y46F (PTP1B TM) or FLAG-PTP1B-D181A-Y46F d406-435 (PTP1B tr. TM) substrate trapping mutants. **D.** Quantification of the calcium response by the determination of the baseline-corrected area under the curve of the calcium kinetics. Statistical analysis was performed using a student’s t-test (two-sided, n=3, error bars = SEM). AUC, area under the curve

**Supplemental Figure 7.**
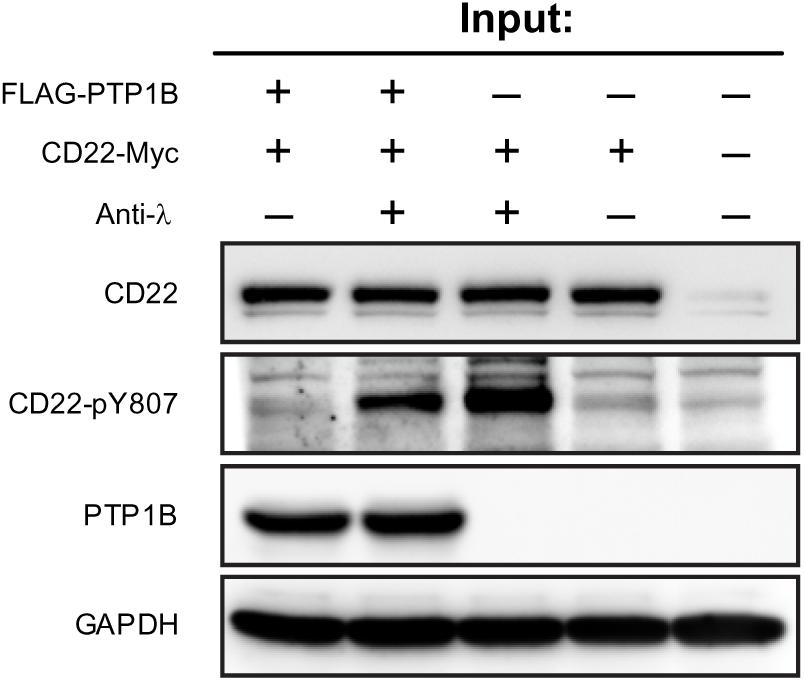
CD22 phosphatase assay. Related to Figure 6 **A.** Lysates used as an input for the CD22 phosphatase assay. As indicated cells were stimulated with 10 µg/ml anti-λ antibody for 5 minutes after 30 minutes of serum starvation. The expression of FLAG-PTP1B was induced by the addition of doxycline to the cell culture media.

## SUPPLEMENTAL TABLES

Supplemental Table 1: The Ramos B cell proteome at an absolute quantitative scale

Supplemental Table 2: Result of the GO Term and REACTOME Pathway enrichment

Supplemental Table 3: Comparison of the copy numbers in Ramos B cells to naïve B cells from Rieckmann et al. 2017

Supplemental Table 4: RNA sequencing data

Supplemental Table 5: GO terms of non-functional transcripts

Supplemental Table 6: Matched proteome and RNA sequencing data

Supplemental Table 7: Copy numbers and TPM values of the classical protein tyrosine phosphatases

Supplemental Table 8: SILAC-based quantitative proteome analysis of PTP1B knockout cells vs. Ramos B cells

Supplemental Table 9: Results of the PTP1B Co-IPs

Supplemental Table 10: Plasmids and Peptides

